# Metabolite-centric Knowledge Graph Unveils Predictive and Mechanistic Insights through Self-supervised Multimodal Learning

**DOI:** 10.64898/2026.10.13.758464

**Authors:** Yuxing Lu, Xinru Guan, Wei Wu, Li Quan, Lin Wang, Li Chen, Jinzhuo Wang

## Abstract

Metabolomics research is advancing toward a comprehensive view of metabolites and their biological functions, yet conventional, dictionary-like databases fall short of presenting these complexities. To overcome this limitation, we developed MetaKG, a metabolite-centric knowledge graph that integrates diverse chemical, biological, and semantic data, coupled with a global self-supervised multimodal learning framework that generates advanced embeddings (MetaKGE) for both predictive and mechanistic analyses. MetaKG outperforms existing models in standard representation learning metrics and across 12 external datasets. Furthermore, MetaKG refines existing database information by detecting misannotated labels and uncovering previously overlooked biochemical relationships. Its predictive capability naturally yields testable biochemical hypotheses, as demonstrated in studies of G6PD deficiency and colorectal cancer. By transforming passive, database-scale information into extractable knowledge and actionable insights, MetaKG paves the way for hypothesis-driven discovery in metabolomics and holds promise for informing future research in clinical biomarker identification and therapeutic development.

## Introduction

Recent advances in metabolomics have deepened our understanding of the roles of small molecules in cellular physiology and homeostasis [1, 2]. Given that metabolites are chemical entities whose structure, reactivity, and interactions are intrinsically tied to their biological functions, deciphering the vast, disjointed data remains a significant challenge. This gap limits our ability to uncover the intricate relationships that govern metabolic processes [3, 4].

Progress in metabolomics owes much to foundational resources [5, 6]. Databases such as KEGG [7, 8], HMDB [9, 10], ChEBI [11], SMPDB [12, 13], and Reactome [14] catalog metabolite identities, properties, and pathways, while tools like MetaboAnalyst [15] and Metaboverse [16] offer platforms for statistical analysis. Although these resources have considerably advanced the field, their dictionary-like information retrieval and data-driven statistical approaches have limited capability in predicting unseen relationships or revealing hidden biochemical connections [17, 18].

There is a pressing need for tools that can integrate diverse data types and employ predictive modeling to meet the current demands of metabolomics advancement.

Knowledge graphs (KGs) and embedding-based methodologies have emerged as powerful, knowledge-driven frameworks in biomedical research [19-28], as demonstrated by applications in drug discovery (e.g., PharmKG [22]) and integrative analysis (e.g., Bioteque [23]). However, metabolomics poses unique demands: a tool must not only map biological relationships but also encode the chemical complexity of metabolites like reactivity, structural diversity, and context-specific functions [29-31]. Existing KGs lack metabolite-specific frameworks tailored to these demands, creating an unmet need for resources that can predict pathways, uncover hidden biochemical mechanisms, and generate actionable biological insights.

To address these limitations, we present MetaKG, a metabolite-centric knowledge graph built on over 2.5 million entities and 23.4 million relationships that integrates diverse chemical, biological, and semantic data into a unified framework. Building on this, we employ a self-supervised multimodal learning approach that aligns chemical properties, biological concepts, and semantic descriptions of metabolites to generate advanced embeddings - MetaKGE. MetaKGE not only outperforms traditional embedding models in predicting metabolite properties but also enables MetaKG to correct misannotations, quantify overlooked pathway interactions, and generate biochemical hypotheses. By open-sourcing the knowledge graph construction process and detailing our embedding-based methodologies, we aim to foster tools that transform passive, fragmented database information into an interactive, knowledge-driven resource that complements and corroborates existing data-driven metabolomics approaches. Together, these advances are expected to refine our understanding of metabolic pathways and promote future biomedical research in biomarker discovery and therapeutic development.

## Results

### Overview of the Metabolite-Centric Knowledge Graph

We constructed MetaKG, a metabolite-centric knowledge graph that unifies fragmented metabolomics data into a single resource for biochemical exploration **(Figure 1a, Supplementary Figure 1)**.

**Figure 1.**
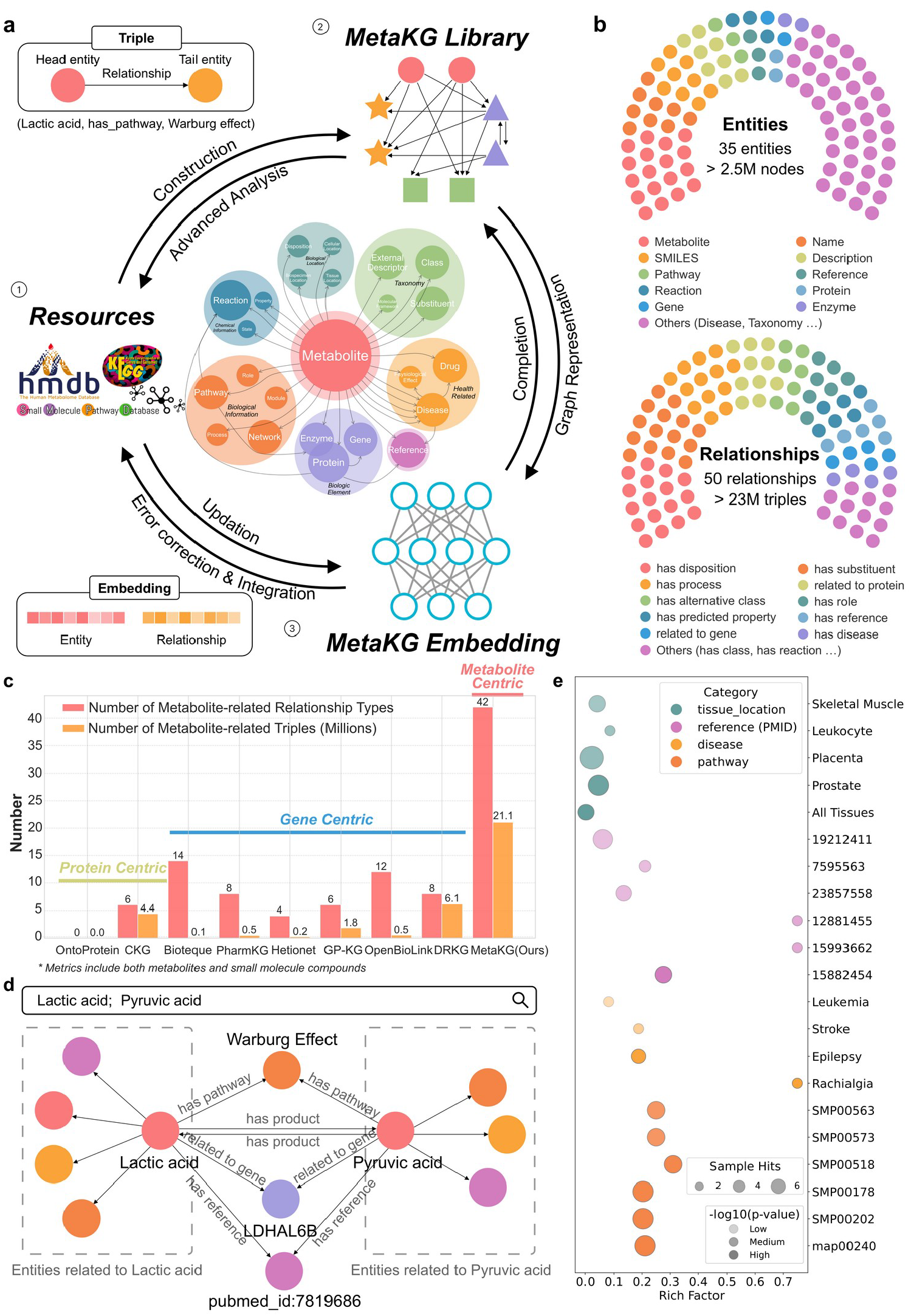
Overview of MetaKG: A Unified, Metabolite-Centric Knowledge Graph. **a)** Knowledge in the database is parsed, organized and constructed into MetaKG, and then represented through learning to obtain MetaKG Embedding. **b)** Distribution of entities and relationships in MetaKG. **c)** As a metabolite-centric knowledge graph, MetaKG has more types of relationships and a larger number of triples related to metabolites compared to other biomedical knowledge graphs. **d)** MetaKG allows for visual representation, showing common or specific relationships between entities. **e)** MetaKG enables enrichment analysis over multi-modal relationships.

Unlike traditional databases that separately catalogue metabolites and pathways, MetaKG integrates multimodal data into a structured network of 2.5 million entities and 23.4 million triples, spanning 35 entity categories (e.g., metabolites, enzymes, diseases) and 50 relationship types. This design captures diverse metabolomics knowledge at unprecedented scale, thereby allowing for more integrated analyses (**Figure 1b, Supplementary Figure 2, Supplementary Table 1,2**).

**Figure 2.**
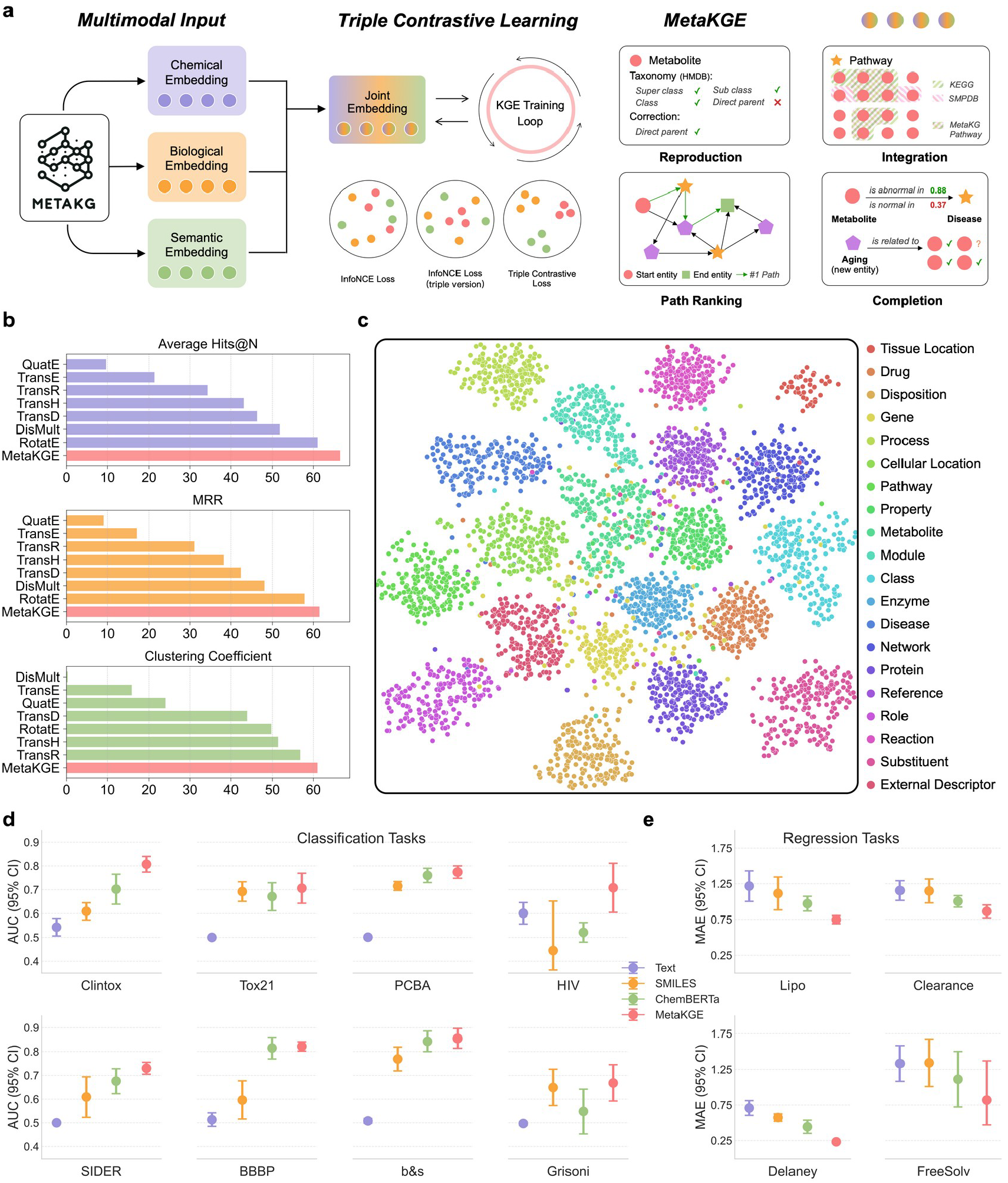
Triple Contrastive Learning Yields Superior MetaKG Embeddings (MetaKGE) for Quantitative and Predictive Analysis. **a)** Overview of the MetaKGE training process. Chemical, biological and semantic representations are integrated and processed through triple contrastive learning to obtain MetaKG embedding. **b)** Comparison of MetaKGE with other knowledge graph representation methods. **c)** 2D UMAP visualization of MetaKGE, where different colors represent different categories of entities. **d)** Performance of MetaKGE on eight external classification tasks, where high area under the curve (AUC) values indicate better performance. **e)** Performance of MetaKGE on four external regression tasks, where low mean absolute error (MAE) values indicate better performance.

Specifically, MetaKG consolidates data from leading metabolomics repositories, including HMDB [9, 10], KEGG [7, 8], and SMPDB [12, 13], and cross-references them with external resources such as PubChem [32], UniProt [33] and ChEBI [11] **(Supplementary Note 1)**. Through a rigorous data curation and alignment pipeline, we transform unstructured data into triples, eliminate redundancies, and connect entities using HMDB IDs as unique identifiers **(Supplementary Figure 3, Supplementary Note 2)**. As a result, MetaKG offers 3-to 10-fold more abundant metabolite-related entities and triples than other knowledge graph tools, positioning it as one of the most extensive metabolite-centric knowledge graphs currently available **(Figure 1c, Supplementary Figure 4)**.

**Figure 3.**
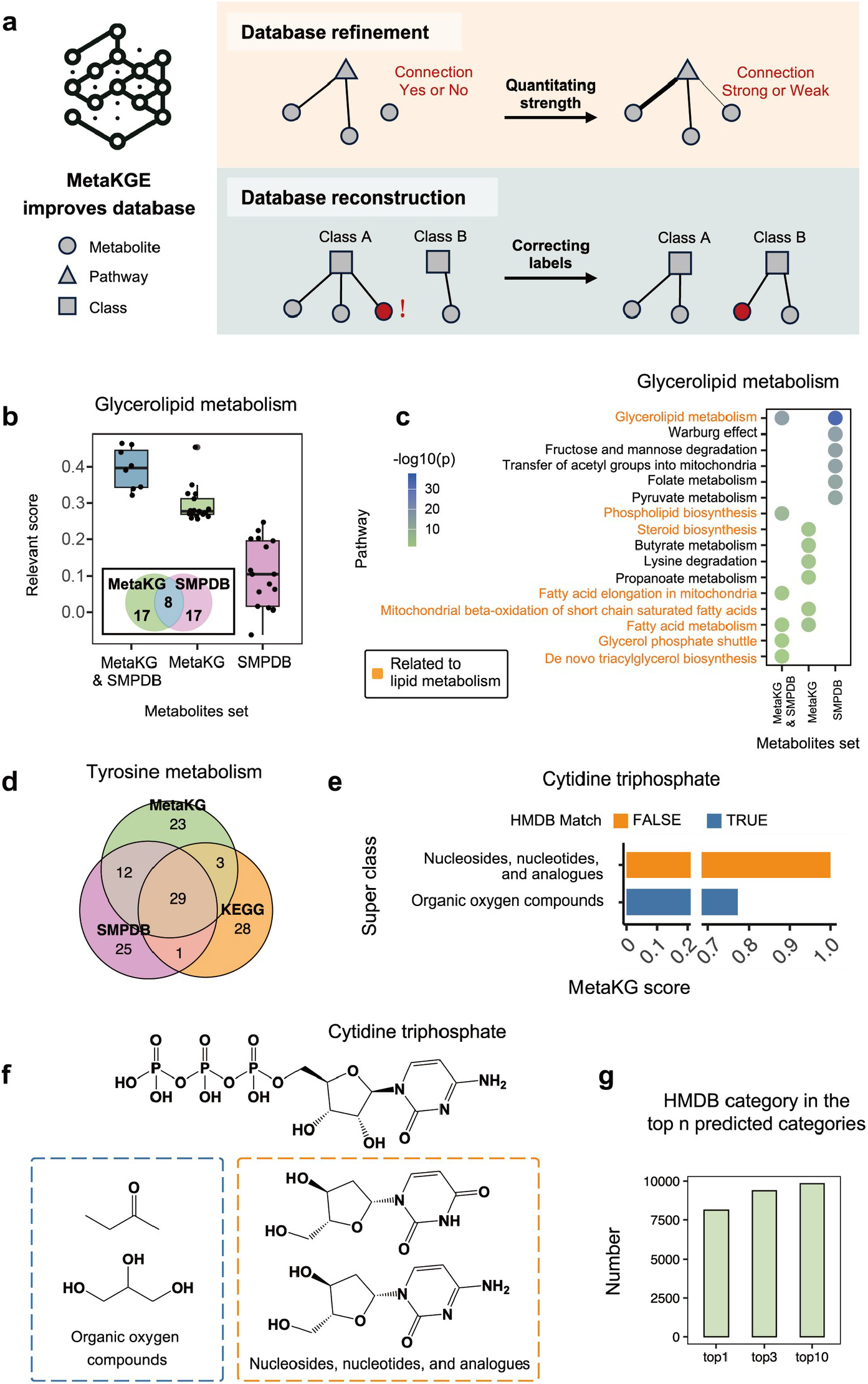
Enhancing Metabolomics Knowledge Through Database Refinement and Reconstruction with MetaKGE. **a)** Diagrams illustrating how MetaKG refines and reconstructs database by quantitating link strengths and correcting labels. **b)** The box plots of metabolite relevance scores for different groups of metabolites. **c)** The enrichment p-values for respective metabolite sets against all pathways in SMPDB. A Venn diagram of the predicted tyrosine metabolism metabolite set from MetaKGE and the record metabolite sets from KEGG and SMPDB. **e)** Prediction of the superclass taxonomic labels for Cytidine triphosphate. **f)** Chemical structures of Cytidine triphosphate and representative metabolites labeled as “Organic oxygen compounds” or “Nucleosides, nucleotides, and analogues”. **g)** The number of top ‘n’ predicted labels that match recorded taxonomic labels, based on predictions for 10000 metabolites.

**Figure 4.**
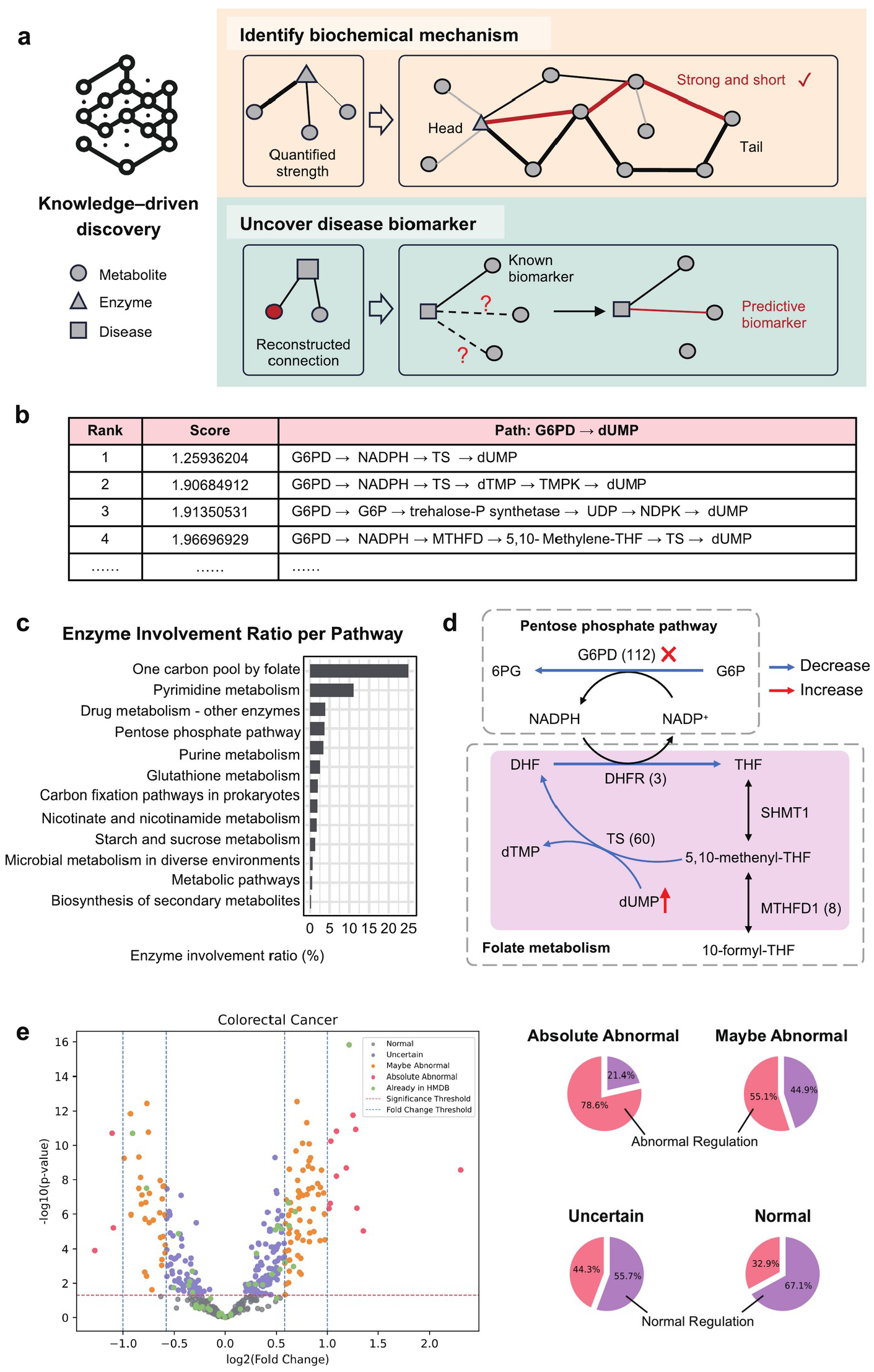
MetaKG links biochemical mechanisms and disease-associated metabolites. **a)** Knowledge-driven discovery framework. **b)** Ranked paths linking G6PD to dUMP. **c)** Enzyme involvement ratios across pathways represented in the inferred paths. **d)** Biochemical connections between the pentose phosphate and folate pathways; numbers in parentheses indicate enzyme occurrences in the analyzed paths. **e)** Colorectal cancer volcano plot and category-specific predictions of normal or abnormal concentration relationships. G6PD metabolomics data were reanalyzed from the published study in reference 41.

This framework supports intuitive exploration tools that go beyond static record retrieval or standard analysis. Researchers can, for example, query all entities related to lactic acid and pyruvic acid (key metabolites in energy metabolism) and visualize the resulting subgraphs **(Figure 1d, Supplementary Figure 5**). MetaKG also extracts entities from sentences to map internal associations and expand to other knowledge in KG **(Supplementary Figure 6)**. Additionally, enrichment analyses and Sankey plots illustrate the flow of metabolomics data across diverse contexts, such as pathway, taxonomy, disease, research articles as well as any other connected entities, offering a comprehensive view for these enriched metabolites **(Figure 1e, Extended Figure 1)**.

**Figure 5.**
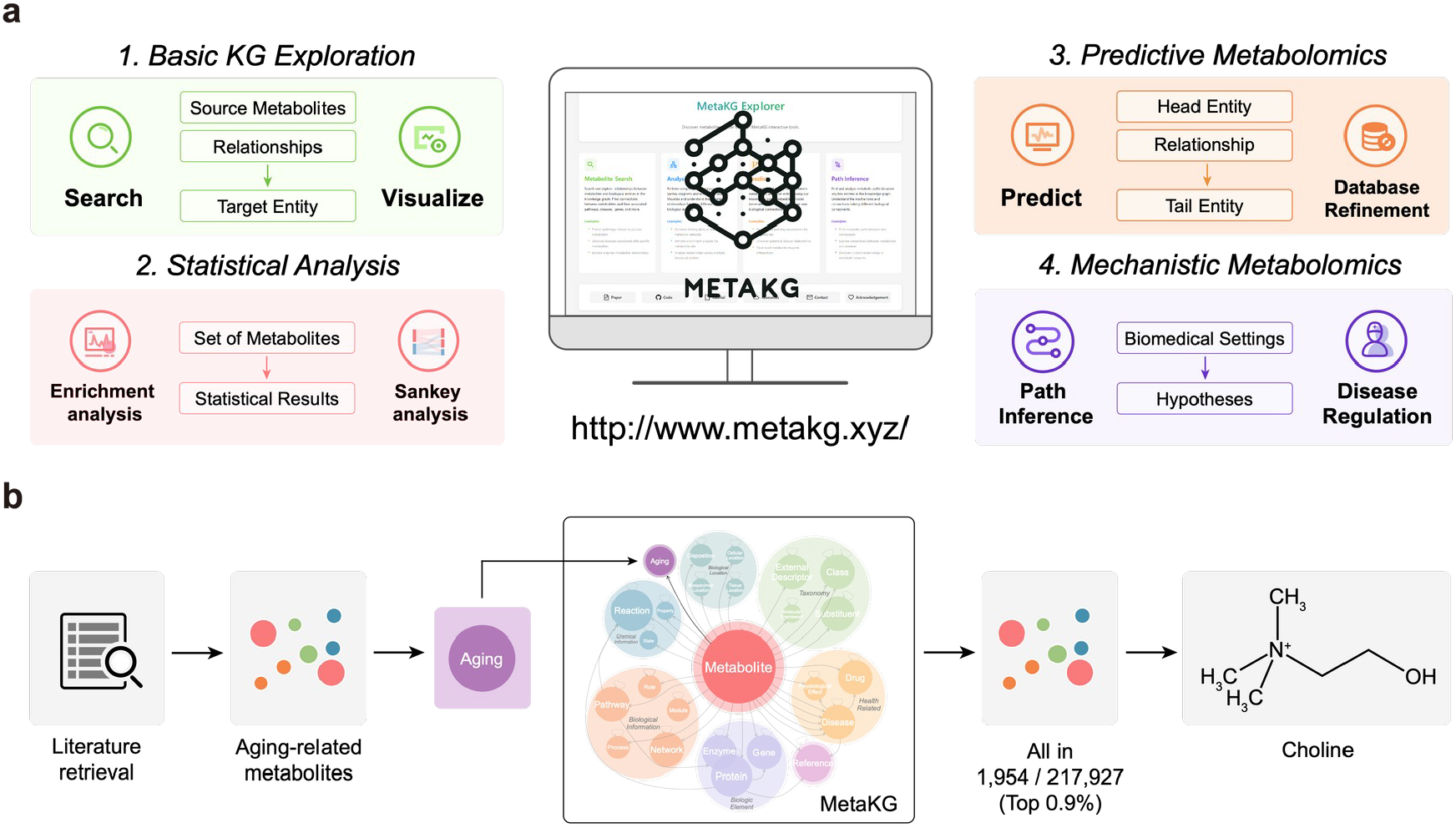
MetaKG as an Open and Evolving Resource for Advancing Metabolomics Research. **a)** MetaKG demonstration website offers users a range of features, including KG exploration, statistical analysis, predictive and mechanistic applications. **b)** By integrating unrecorded knowledge, MetaKG can automatically update itself through retraining, enabling it to conduct analyses and make predictions in new domains.

By forging a multimodal, interconnected foundation from diverse sources, MetaKG transcends the limitations of dictionary-like databases and prepares metabolomics information for predictive modeling.

### Predictive Embeddings through Self-supervised Multimodal Integration

MetaKGE unlocks MetaKG’s predictive potential by converting its 23.4 million triples into high-dimensional embeddings. By integrating chemical structures, biological roles, and textual descriptions, MetaKGE drives predictive tasks such as KG completion, pathway relevance scoring, and biochemical hypothesis generation, surpassing the limitations of single-modal models **(Figure 2a)**.

We trained MetaKGE using self-supervised knowledge graph embedding framework with a triple contrastive learning module that aligns information from different modalities into a unified embedding space **(Supplementary Figure 7, Supplementary Note 3)**. Specifically, ChemBERTa-2 [34] encodes molecular features from chemical structures; SciBERT [35] processes textual descriptions into semantic vectors; and RotatE models MetaKG’s relational structure, preserving biological connections. Synchronizing these modalities yields embeddings that integrate diverse metabolite properties with enhanced predictive power.

Compared with widely used knowledge graph embedding approaches, including TransE [36], TransR [37] and RotatE [38], MetaKGE achieved the highest Hits@1, Hits@3, Hits@5, Hits@10, mean reciprocal rank (MRR) and clustering coefficient among the models in Supplementary Table 3. Hits@10 improved by 12.7% relative to RotatE. Mean rank (MR) improved from 895 for RotatE to 643 for MetaKGE, although TransD achieved a lower MR of 578 (Figure 2b). UMAP projections [39] illustrate grouping of related entities, including “acute liver disease” and “acute liver failure” (Figure 2c, Extended Figure 2), and tissue-related groupings (Supplementary Figure 8).

In head-to-head comparisons on eight external classification tasks (ClinTox, Tox21, PCBA, HIV, SIDER, BBBP, b&s, Grisoni) and four external regression tasks (Lipo, Clearance, Delaney, FreeSolv) **(Supplementary Table 4, Supplementary Figure 9)**, MetaKGE outperforms single-modal pretrained embeddings (SciBERT [35], Mol2Vec [40] and ChemBERTa-2 [34]), achieving 0.86-17.80% higher AUROC values and 14.04-47.75% lower MAEs **(Figure 2d,e)**.

Collectively, by providing an advanced triple contrastive learning framework that synergizes chemical, semantic, and relational data, MetaKGE demonstrates its superiority in metabolite embeddings, setting the stage for the quantitative and predictive metabolomics analyses.

### Correcting and Quantifying Metabolomics Knowledge with MetaKGE

Metabolomics thrives on reliable data, yet inconsistencies often plague existing databases. Rather than serving as a static repository, MetaKG leverages the quantitative embeddings of MetaKGE to actively refine and reconstruct existing metabolomics knowledge, addressing data gaps and misannotations (**Figure 3a**).

Most relational links between a metabolite and other entities are binary - yes or no - with no granular measure of the connection strength. MetaKG fills this gap with a quantitative framework that evaluates the metabolite relevance within metabolic pathways. For example, metabolites with high relevance scores in the glycerolipid metabolism pathway were enriched in pathways related to lipid and fatty acid metabolism, whereas lower-scoring metabolites showed less associations **(Figure 3b,c)**. This nuanced approach not only yields a more accurate and granular representation of metabolic pathways, but also highlights disparities among sources. For example, less than 45% of metabolites in the tyrosine metabolism pathway overlap between the SMPDB and KEGG databases. MetaKG bridges these gaps by generating a unified list of high relevance score metabolites that better reflects biological relationships across diverse resources **(Figure 3d, Supplementary Figure 10, Supplementary Table 5)**.

MetaKG’s quantitative framework also enables systematic database reconstruction. A key advantage is to detect the inconsistencies arising from manual curation or algorithmic pipelines. By assigning a confidence score to each metabolite’s taxonomic label, MetaKG ensures that high-confidence labels align well with existing curated entries. Notably, MetaKG reclassified Cytidine triphosphate (HMDB0000082) from the existing super class label “Organic oxygen compounds” to the more accurate “Nucleosides, nucleotides, and analogues” **(Figure 3e)**. This correction was supported by both structural comparison and the description in the database **(Figure 3f)**. In a broader analysis of 10,000 metabolites, MetaKG flagged metabolites with potential taxonomic inconsistencies **(Figure 3g, Supplementary Table 6)**, suggesting such large-scale checks can efficiently detect potential inconsistency for refinement in static repositories.

By integrating quantitative relationship scoring with active knowledge reconstruction, MetaKG enhances the depth and accuracy of current metabolomics information. In turn, these refinements strengthen MetaKG’s foundational data, forming a virtuous cycle where refined databases and robust embeddings reinforce each other, transforming how metabolomics information is trusted and utilized.

### Knowledge-Driven Insights into Biochemical Mechanisms and Disease Signatures

Data-driven analysis tools can spot patterns, but their reach is often confined by experimental scope. MetaKG introduces a knowledge-driven framework that enhances and expands the capabilities of data-driven metabolomics tools. By integrating diverse multimodal data, MetaKG provides a more comprehensive perspective, revealing hidden relationships and biological mechanisms that might otherwise remain obscured **(Figure 4a, Supplementary Note 4)**.

An example of this synergy is MetaKG’s application to biochemical mechanism inference in a G6PD knockout model [41]. Glucose-6-phosphate dehydrogenase (G6PD) plays a critical role in generating NADPH for downstream metabolic processes. Studies have reported a ∼100-fold increase in dUMP levels in G6PD knockout cells **(Supplementary Figure 11, Supplementary Table 7,8)**. MetaKG identified potential pathways from G6PD to dUMP. After statistically analyzing the pathway attribution of all enzymes in the top 20 paths, it was discovered that enzymes in the “one carbon pool by folate” pathway appeared most frequently.

By querying all metabolic pathways linking G6PD to dUMP and applying quantitative metrics, MetaKG identified key intermediates such as NADPH and thymidylate synthase, aligning with the reported biochemical mechanisms **(Figure 4b,c, Supplementary Table 9)**. Further statistical analysis of the frequency of all enzymes in these paths revealed that G6PD and thymidylate synthase (TS) were the two most frequently occurring enzymes. This indicates that in the context of a G6PD knockout, there may be a metabolic process affecting dUMP with G6PD and TS as key nodes. **(Figure 4d, Supplementary Figure 11, Supplementary Table 10,11)**. Importantly, MetaKG identified these mechanistic intermediates without relying on experimental data, demonstrating its unique strength as a knowledge-driven framework for reliable hypothesis generation.

In disease contexts, MetaKG enhances the predictive power of data-driven tools by uncovering hidden metabolic relationships. For instance, in a colorectal cancer cohort (n=31, **Supplementary Table 12**), MetaKG identified 12 of 15 (80%) dysregulated metabolites as having “abnormal concentrations”, whereas 41.4% (182 of 440) of the remaining metabolites were classified as abnormal **(Figure 4e, Supplementary Table 13)**. These findings indicate that even in the complex disease context, MetaKG’s knowledge-driven approach can yield valuable hypotheses for further biomarker investigation.

In summary, MetaKG generates valuable hypotheses for data-driven experiments. By drawing on a quantitative understanding of the entire metabolite-centric knowledge graph, it provides a broad biological context that complements the interpretation of experimental findings. This synergy between knowledge-driven insights and experimental data accelerates discovery in metabolomics research by enhancing hypothesis generation.

### Advancing Metabolomics Research through FAIR and Usable Resources

MetaKG is an open platform designed to support FAIR principles that empowers metabolomics research with interactive exploration and predictive insights [42] **(Supplementary Note 5)**.

Accessible at http://www.metakg.xyz, its demonstration website showcases basic KG exploration, statistical analysis, predictive metabolomics and mechanistic metabolomics, which are built on MetaKG’s refined knowledge base and advanced embeddings **(Figure 5a, Supplementary Note 6)**. For example, researchers can perform enrichment analyses to identify multimodal metabolic patterns or use inference tools in a user-friendly workflow that brings mechanistic hypotheses within reach.

The MetaKG has been open-sourced at https://github.com/YuxingLu613/MetaKG, alongside tutorials and reproducibility reports. The codebase encompasses the entire MetaKG pipeline, including data parsing, knowledge graph construction, data exploration and statistical analysis, MetaKGE representation learning, and advanced inference applications **(Supplementary Figure 12)**, providing users with the flexibility to adapt the platform to their own data and tasks. Meanwhile, all resources, including the preprocessed data and MetaKG embeddings, are released publicly and will be updated regularly to reflect newly integrated datasets and algorithmic improvements.

MetaKG is also extensible, featuring external links to widely used biological databases (e.g., PubMed, ChEBI, and UniProt) and the capacity to integrate custom datasets, relationships, or additional omics layers. Users can adapt MetaKG to emerging scientific questions or specialized applications by contributing private knowledge to expand the graph and develop new prediction tasks. For instance, we curated a dataset of 235 aging-related metabolites (Supplementary Table 14). Five metabolites with high identifiers and sparse known information were excluded; 180 metabolites were incorporated into MetaKG for training, and 50 were reserved for testing. After retraining MetaKGE, the model ranked the 50 test metabolites among the top 0.9% in relevance scores **(Figure 5b)**. One high-ranking hit, choline, though not covered in the 235 metabolites, is linked to longevity pathways [43-45], suggesting MetaKG has already unlocked its predictive capabilities in the context of aging, an area previously uncovered in the original KG. This flexibility ensures that MetaKG evolves with new datasets and shifting research priorities.

In summary, MetaKG establishes an open, reproducible, and extensible knowledge graph framework for the metabolomics community. By synergizing its predictive and mechanistic metabolomics capabilities with existing metabolomics databases and analytical tools, MetaKG enables deeper insights into metabolic processes and accelerates innovative applications in biomarker discovery, therapeutic development, and beyond.

## Discussion

Recent advances in metabolomics have exposed the intrinsic limitations of dictionary-like databases and catalogued pathway maps, which struggle to cross-reference and connect the vast number of metabolites and associated chemical, biological, and semantic information. To address these challenges, we developed MetaKG, the largest metabolite-centric knowledge graph to date, by unifying 2.5 million entities and 23.4 million triples from repositories such as HMDB, KEGG, and SMPDB. The interconnected nature of MetaKG is demonstrated by integrative analyses across diverse domains, including metabolite enrichment analysis spanning pathways, taxonomy, diseases, and research literature **(Figure 1)**. Building on this unified framework, we employ a triple contrastive learning approach to integrate multimodal data into unified quantitative embeddings (MetaKGE).

These embeddings outperform other published embedding methods (e.g., TransH and RotatE), as evidenced by superior performance in eight external classification tasks and four regression tasks **(Figure 2)**. By capturing the diverse yet nuanced relationships among metabolites, their chemical properties, and their biological interactions, MetaKG and MetaKGE open the door to quantitative understanding and predictive modeling of metabolomics knowledge **(Extended Figure 3, Supplementary Table 15)**.

Conventional metabolomics databases are typically viewed as ground-truth sources for biological insights, yet curation errors and inconsistencies across databases remain largely unaddressed. MetaKG bridges this gap by introducing a quantitative perspective that re-examines relationships, such as confidence scores for metabolite taxonomic classifications and relevance scores for metabolite– pathway connections, thereby enhancing the robustness of curated data (**Figure 3**). Moreover, MetaKG complements data-driven analysis tools like MetaboAnalyst and Metaboverse. For instance, in a G6PD knockout study where dUMP metabolite accumulates, MetaKG’s identification of mechanistic intermediates linking G6PD and dUMP stems from its quantitative, predictive understanding of the enzyme-metabolite connection instead of experimental measurement data (**Figure 4**). Such knowledge-driven hypothesis, validated in this case by experimental data, can further enhance data-driven analysis results to yield deeper insights in future studies. Together, these examples highlight MetaKG not only provides high-quality knowledge-driven hypothesis, but also can cooperate with other large-scale metabolomics resources to refine each other.

MetaKG’s demonstration website showcases its interconnected contents and predictive modeling, from enrichment analyses to mechanistic pathway inference, within a user-friendly workflow. The platform is fully open-sourced, providing reproducible codes and preprocessed data that users can adapt for custom development. While MetaKG can already incorporate manually curated information **(Figure 5)**, we envision expanding its capabilities through MetaKG-LLM, which uses instruction-based prompts and generative models GPT and DeepSeek to extract knowledge triplets from new literature [46-48], thus automating the flow of information from curation to integration into MetaKG. The MetaKG Family toolkit, which includes the MetaKG construction phase, the MetaKGE embedding phase, and the proposed MetaKG-LLM automation phase, ensures that MetaKG continue to evolve with the metabolomics and AI communities **(Extended Figure 4)**.

Admittedly, limitations and challenges remain. On the data side, MetaKG’s accuracy is fundamentally dependent on the quality and scope of its source databases like HMDB and KEGG. Since these repositories contain inconsistencies and curation errors, these are inevitably propagated into the knowledge graph. Unifying these disparate sources also presents challenges; standardizing metabolites to a HMDB ID is applied in MetaKG but this reliance on existing cross-reference mappings can lead to an incomplete representation by excluding metabolites not found in HMDB. Furthermore, on the metabolomics side, representing the diversity of biological concepts introduces complexity. For instance, while pathway maps emphasize direct biochemical connections, a pathway analysis might need to include indirect regulators such as allosteric or signaling molecules. Therefore, further expansion of entity and triple types in MetaKG is essential to capture these subtle contextual differences. From the AI perspective, the framework has several constraints. The high-dimensional embeddings generally lack direct biological interpretability and may underperform for understudied metabolites. While MetaKG is extensible and can incorporate curated datasets through retraining, its architecture is not designed for the direct, on-the-fly statistical analysis of raw experimental data in the manner of tools like MetaboAnalyst. Future refinements could leverage attention-based architectures or predefined latent spaces to improve both predictive performance and interpretability [49-51]. Much like early large language models, which occasionally display errors and biases, we view these imperfections as opportunities for continued improvement.

In conclusion, MetaKG marks an early yet transformative milestone in knowledge-driven metabolomics analysis. By integrating a quantitative, interconnected knowledge graph with predictive and mechanistic insights, MetaKG offers an open, evolving framework ready to serve and grow with the metabolomics community. We envision that a collaborative ecosystem, where metabolomics databases, data-driven analytical tools, and knowledge-driven discovery methods synergize and co-evolve, will usher in a new paradigm for the metabolomics research, ultimately driving mechanistic breakthroughs in biomedical innovation.

## Methods

### Data sources for MetaKG

MetaKG is an extensive knowledge graph that consolidates information from a diverse array of online metabolomics databases, providing users with a comprehensive and multidimensional resource for metabolomics research. It sourced data from reputable platforms such as HMDB (https://hmdb.ca/) [9], KEGG (https://www.genome.jp/kegg/) [8], SMPDB (http://smpdb.ca/) [12] and leaves cross-reference links for PubChem (https://pubchem.ncbi.nlm.nih.gov/) [32], CheBI (https://www.ebi.ac.uk/chebi/) [11], FooDB (http://foodb.ca/) [52], and UniProt [33], etc. **(Supplementary Table 1,2)**. These databases, which have been the subject of numerous research studies and have gained a reputation for their meticulous curation over time, provided a wealth of information on metabolites, including chemical properties, biological roles, and pathway associations [53, 54].

### Data preparation for MetaKG

We developed a multi-stage data preparation workflow to construct MetaKG from heterogeneous sources such as HMDB, KEGG, and SMPDB **(Supplementary Figure 3, Supplementary Note 2)**. Our process began with data acquisition and parsing, where we tailored custom parsers for each database’s distinct file formats. Each extracted data element was standardized into a unified representation: entities were stored as lists in the format {entity name, category}, and relationships were encoded as triples {head entity, relation, tail entity}. Following extraction, the next stage involved knowledge cleaning. We implemented automated rule-based filtering to identify and remove overly generic or erroneous associations. For instance, associations with non-specific descriptions such as “Health effect” were discarded. These types of entities often have a very large degree, meaning they are linked to a vast number of other entities across MetaKG, making them highly ambiguous and not informative. We then leveraged external cross-reference mappings from HMDB, KEGG, and SMPDB to align entities with disparate identifiers. Entities representing the same biochemical concept were interconnected via explicit “the_same_as” relationships. Additionally, we standardized entity names by normalizing letter casing and removing special symbols to minimize duplicate entries and facilitate matching across databases. Similar relationships were merged into a single standardized term to ensure clarity and consistency. Quality control measures were applied to identify and handle missing numeric values and outliers using statistical methods. Baseline metabolite concentrations in healthy individuals were established and then compared with patient data to construct relationships that indicated upregulation or downregulation in disease contexts. Metabolites and related entities were standardized to HMDB IDs, ensuring a unique and consistent representation across the graph. Additionally, similar or synonymous relationships were consolidated to eliminate redundancy, resulting in a coherent dataset that underpins MetaKG.

We employed *Neo4j* (http://neo4j.com/) [55] as a tool for knowledge graph management and visualization. *Neo4j* is a widely-used graph management tool that represents knowledge graphs as graph structures consisting of nodes and relationships. By utilizing the Cypher query language in Neo4j, we can create and define node and relationship patterns. *Neo4j* offers built-in graph algorithm libraries, enabling advanced querying and analysis tasks like path analysis and similarity calculations. Users can also make use of *Neo4j*’s visualization tools to explore knowledge graph data in an intuitive manner (**Supplementary Figure 13**). Moreover, we utilized the *networkx* package (https://networkx.org/) [56] to develop functions for statistical analysis and path searching.

### Multimodal embedding

We employed a multi-modal embedding approach that integrates both chemical and textual information. For each metabolite, we retrieved its corresponding SMILES sequence and textual description from MetaKG. Specifically, for each *m⊆ M*, where *M* is the set of all metabolites in MetaKG *G*, we used the MetaKG’s search function to identify the corresponding SMILES sequence *s⊆S* and text description *t ⊆T* . Here, *S* is the set of all SMILES sequences in *G*, and *T* is the set of all text descriptions in *G*. The SMILES sequences were encoded using the ChemBERTa-2 [34] model (https://huggingface.co/DeepChem/ChemBERTa-77M-MLM), and the global information for each metabolite was extracted from the final hidden state of the [CLS] token, yielding a vector *m*_*s*_ *∈* ℝ^768^ ; Similarly, the textual descriptions were encoded using the SciBERT [35] model (https://huggingface.co/allenai/scibert_scivocab_uncased), with the corresponding global information represented as *m*_*t*_ *∈* ℝ^768^ from the [CLS] token head. These encodings were used to generate rich, multi-modal embeddings that capture diverse data formats. To facilitate integration into the MetaKGE training framework, the 768-dimensional vectors *m*_*s*_ and *m*_*t*_ underwent average pooling operations to align their dimensions with those of the MetaKG embeddings *m*_*emb*_ *∈* ℝ _128_, ensuring compatibility for further incorporation into the model training process.

### MetaKGE Algorithm

The MetaKGE Algorithm (Pseudo-code in **Supplementary Figure 7**) optimized the embeddings for entities and relationships in the MetaKG *G* through two training phases: 1) multimodal training and 2) combined training. In the multimodal training phase, for each epoch, we iterated over batches of triples ( *s, p, o* ) and computed the score for each triple, here we employed RotatE [38] as an example.

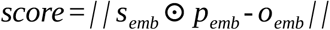

where *s*_*emb*_, *p*_*emb*_, and *o*_*emb*_ are the embedding of the head entity *s*, relationship *p*, and tail entity *o* respectively. *⊙* represents for Hadamard product, which can be regarded as rotating the entity embedding *s*_*emb*_ by the relation embedding *p*_*emb*_. The batch loss *L* is the sum of these scores, and embedding metrics for entities *E∈* ℝ^( *num* _ *entities ×d* )^ and relationships *R∈* ℝ^( *num* _ *relations ×d* )^ were updated via gradient descent. To facilitate multimodal integration, we update each metabolite’s embedding *m*_*emb*_ using its SMILES-based chemical information vector *m*_*s*_ and text-description-based semantic information vector *m*_*t*_ through a triple contrastive learning module to learn an integrated vector for multiple modalities.

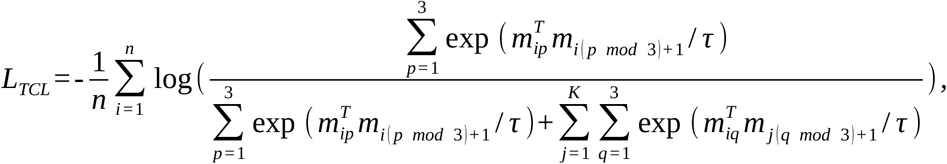

where *n* is the number of samples, *τ* is a temperature parameter, *m*_1_, *m*_2_, *m*_3_ represents *m*_*emb*_, *m*_*s*_ and *m*_*t*_ respectively. *K* is the number of negative samples. We then update *m*_*emb*_ as the integrated multimodal representation for metabolite *m* by calculating the average vector

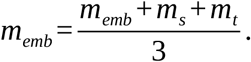

In the first phase, all multimodal representations were trained together in every training loop with triple contrastive learning [57], ensuring a deep-level alignment of these different information. While in the combined training phase, the process was similar to the multimodal training phase but was focused solely on optimizing *E* and *R* based on *G* to attain a more robust and more structural representation, without updating the multimodal embeddings. In essence, the MetaKGE algorithm presented a structured procedure to collaboratively optimize entity and relationship embeddings while seamlessly integrating multimodal information, ensuring a richer representational power.

### MetaKGE training settings

MetaKGE training was carried out using triples in MetaKG. This dataset was meticulously partitioned into training, validation, and testing sets using an 8:1:1 ratio. The choice of hyperparameters was designed to maximize both efficiency and reliability of the model. Both the entity and relationship embeddings were initialized with 128 dimensions. For optimization, the margin ranking loss

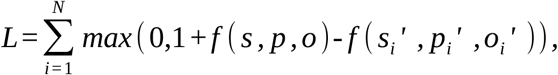

was employed in conjunction with the AdamW [58] optimizer, where *f* ( *s, p, o* ) represents the score computed by the model for a positive triple ( *s, p, o* ) that actually exists in the knowledge graph, and *f* ( *s*_*i*_′, *p*_*i*_′, *o*_*i*_′) is the score for a triple that is assumed not to exist in the knowledge graph. The learning rate was set at 1.0 *×* 10^-3^. The training process spanned 1000 epochs and was executed on a single Tesla A100 GPU. Depending on the foundational KGE model used, the total computation time ranged between 3 and 60 hours. To maintain the reproducibility of our experimental results, we employed a fixed random seed of 42. The selection of these settings was carefully accomplished to establish a balance between computational efficiency and the fidelity of the model, thereby ensuring both robust and reproducible results. When reporting experimental results of MetaKGE, we used common metrics such as Hits@1, Hits@3, Hits@5, Hits@10, MR, MRR, and clustering coefficient to evaluate performance. Hits@n measures the percentage of samples in the test set for which the true target appears within the model’s top *n* predictions.

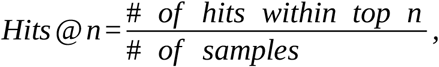

MR (Mean Rank) calculates the average rank of the true targets across all samples in the test set.

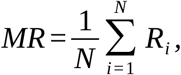

where *R*_*i*_ is the rank of true target in sample *i*. MRR (Mean Reciprocal Rank) computes the average of the reciprocal ranks of the true targets for each sample in the test set.

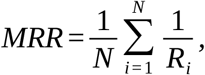

These metrics were widely employed for assessing the performance of models in knowledge graph embedding and information retrieval tasks.

### MetaKG search and subgraph visualization

In both the MetaKG package and our web server, we implemented advanced functions for KG search and subgraph visualization, designed to facilitate user engagement with MetaKG. Formally, MetaKG was represented as *G*={*E, R, T* }, where *E* is the set of entities, representing the set of entities in the graph, *R* denotes the set of relationships or predicates, connecting pairs of entities, and *T* ={( *s, p, o* ) */ s, o′ E, p′ R* } is the set of triples. Each triple ( *s, p, o* ) indicates that there exists a relationship *p* between *s* and *o*.

#### MetaKG search

Querying within the context of a KG involved the identification of a subset *T ′ ⊆T* that aligns with specified criteria. For example, in forward searches, given *s ′* and *p ′*, the goal was to find all *o ′* such that ( *s ′, p ′, o ′* )*′ T* . Conversely, backward searches operated on the opposite principle, starting from *o ‘* to find *s ′*.

#### Subgraph visualization

A subgraph *G′* of MetaKG was defined by *G′* ={*E’, R ′, T ′* }, where *E ′* ={*s /* ( *s, p, o* )*′ T ′* }*∪* {*o /* ( *s, p, o* )*′ T ′* }, and *R ′* ={ *p /* ( *s, p, o* )*∈ T ′* }. Once obtained the subgraph *G′* from either MetaKG search or KG construction from a subset of triples, visualization tools like *Neo4j* and *echarts* used in our website can then be applied to render *G′*, thereby offering users a visual insight into the intricate web of relationships and entities encapsulated within *G′*. This graphical depiction aids in the deeper comprehension and exploration of the underlying data structure, enriching user’s research experience.

### Enrichment analysis

MetaKG facilitated enrichment analysis for various entity types by harnessing their comprehensive multimodal knowledge. In the context of analyzing a set of differentially expressed metabolites, the process began with the collection of corresponding multimodal information from MetaKG. Subsequently, enrichment was quantified using the hypergeometric distribution test method, a statistical approach designed to assess the significance of overlap between two sets. The probability mass function for the hypergeometric distribution is given by:

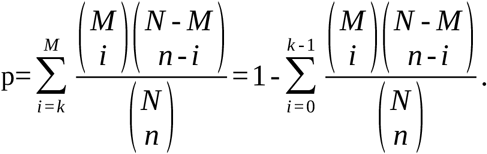

Here, *N* represents the total number of all considered metabolites, *M* is the count of differentially expressed metabolites, *n* stands for the total number of metabolites associated with a specific entity, and *k* denotes the count of differentially expressed metabolites related to that entity. Enrichment analysis of the pathway metabolite set used the hypergeometric distribution algorithm. In **Figure 3**, the pathway of a metabolite set predicted by MetaKG was enriched in pathways of SMPDB or KEGG respectively, while the metabolite set from a specific pathway in SMPDB was enriched in pathways of KEGG or MetaKG. The same applies to KEGG.

### Embedding visualization and evaluation

We utilized the UMAP [39] algorithm to generate a 2D representation of the embeddings. To visualize the complete set of entities (**Figure 2c**), we chose 200 data points from each category. In cases where a category contains fewer than 200 entities, all data points from that category was displayed in the graph. Specifically, for visualizing tissue location entities (**Supplementary Figure 8**), we selected 200 data points from each of the following categories: Fibroblasts, Placenta, Brain, Liver, and Kidney.

In order to evaluate the performance of MetaKGE’s metabolite representations in various biological and chemical tasks, we selected 12 external datasets from different sources like HuggingFace, Kaggle, and GitHub **(Supplementary Table 4)**. We extracted metabolites relevant to human metabolomics from these datasets and split each dataset into training and testing sets using 5-fold cross-validation and selected the SMILES-based and text-based embeddings for comparison. For the classification tasks, we used a deep neural network (**Supplementary Figure 9**) designed based on the skip-connect architecture [59]. For each metabolite’s embedding *m*_*emb*_, the model classified it using

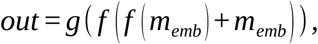

where *g* ( *·*) denotes a classification neural network, and *f* ( *·*) stands for multilayer perceptron (MLP) network. To evaluate the performance, we used micro-AUC for classification tasks and mean absolute error (MAE) and mean squared error (MSE) for regression tasks; Figure 2e reports MAE. For hyperparameter optimization, we employed a grid search to fine-tune key parameters. During hyperparameter tuning, we report the highest performance with mean value and 95% confidence interval (CI) to robustly evaluate model performance across different subsets.

### Metabolite taxonomy prediction

For each metabolite, we predicted its corresponding taxonomy information, including super class, class, sub class, and direct parents. We selected all results related to metabolite classification and identified the top 10 results for statistical analysis. Subsequently, we normalized the prediction results, setting the highest prediction score to 1, with the remaining scores ranging from 0 to 1. We analyze the prediction results for classification of the top 10000 metabolites in HMDB according to HMDB id. Then each metabolite is formatted into triples using the structure

{hmdb_id:HMDB0000001, has_class, ?} to predict the taxonomic tail entity. The metabolites with the highest MetaKG prediction scores, which also matched the HMDB records, were chosen as representative substances for each taxonomic classification. The prediction process does not consider the structure of metabolites but relies solely on the contextual information recorded in the knowledge base. After obtaining the prediction results, the predicted accuracy of misclassified metabolites is manually verified based on the structural information documented in the database.

### Predicting metabolite relevance to pathways

Different metabolites in a pathway may have varying levels of importance, which was not recorded in existing experiments and databases. We used MetaKGE to rank the importance of metabolites within a pathway. Specifically, for each pathway in KEGG and SMPDB, we predicted the importance ranking of the metabolites included in pathway A using the relationship {metabolite X, has_pathway, pathway A}. During analysis, we first extracted all SMPDB and KEGG pathway metabolites and their corresponding scores to these pathways from MetaKGE. For each metabolite, we define a relevance score to a pathway by subtracting the distance score of that pathway to the average distance score of all pathways (833). Based on the relevance score and the number of metabolites in SMPDB pathways, we selected the most relevant metabolites to form the metabolite set predicted by MetaKGE for each pathway. We used the *KEGGREST* package in *R* (version 1.38.0) to obtain the metabolite set for KEGG pathways, matched KEGG metabolites with HMDB metabolites by name, and removed metabolites from KEGG that do not have a corresponding HMDB ID.

### Path finding and distance ranking

Given a start entity and an end entity, MetaKG can find all the paths between them through the Bidirectional Search [60] algorithm which simultaneously performs two searches: one forward from the start entity and one backward from the end entity. This approach reduces the search space and improves efficiency compared to unidirectional search algorithms like Depth-First Search (DFS) [61]. This approach can be mathematically expressed by maintaining two sets of paths *P*_*forward*_ and *P*_*backward*_ representing the paths developed from the start entity and the end entity, respectively. The algorithm iteratively explored each node *v* adjacent to the current node *u* in both directions, using the following recursive relation:

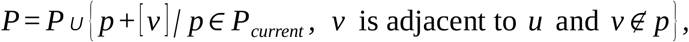

Here, *P*_*current*_ represents the paths developed up to the current node *u*, and *v* denotes appending *v* to path *p*. The algorithm continues to explore nodes in both directions until the forward and backward searches meet at a common node or the number of paths found reaches a predefined maximum limit. We then calculated the total distance for each path based on MetaKGE by summing the reciprocals of the scores for each edge in the path:

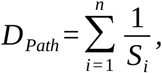

where *n* is the number of edges in the path, and *S*_*i*_ represents the score of the *i*-th edge in the path. We ranked all the paths according to their total distances. We finally merged paths with the same metabolites and enzymes as main entries and sorted them, with paths having different reactions as subentries.

### G6PD knockout model

The metabolite response intensity data for glucose-6-phosphate dehydrogenase (G6PD) knockout in Supplementary Figure 11a were obtained from published literature (PMC6489125). The data were used for peak identification and integration using MAVEN and El-MAVEN software. The heatmap presents the data as the log2 ratio of G6PD knockout to WT.

To explore the ability of data-driven metabolomics analysis methods to elucidate the mechanism by which G6PD KO affects dUMP, the data from HepG2 cells were analyzed using the Global Test in the Pathway Analysis module of MetaboAnalyst 6.0 and the Collapse Reaction function in Metaboverse v0.11.4.

### Metabolite regulation prediction

For each dataset used to calculate differential metabolites, we first performed missing value imputation and normalization on the dataset. Specifically, we used the mean strategy to fill in missing values and employ Min-Max normalization to scale the data, scaling all numerical features to a range of 0 to 1.

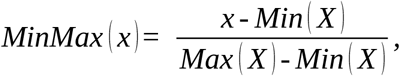

where *x* is the expression value of a metabolite in one sample, and *X* is the expression value list of the cohort. Then we divided the data into disease and normal group, and calculated the fold change of the expression level of each metabolite in the disease group compared to that in the normal group. At the same time, we used the P-value to determine whether the expression level of each metabolite in two groups was significantly different. We considered the P-value threshold of 0.05 and the fold-change threshold of 2, and separate the volcano plot into normal regulated and abnormal regulated areas. In predicting the regulatory relationship between metabolite *m* and disease *d*, we obtained the scores of the two triples {m, has_normal_concentration_in, d} and {m, has_abnormal_concentration_in, d} through the MetaKGE’s score function and then sort them. The prediction with the higher score was selected as the final result. Baseline accuracies were calculated as the proportion of abnormally regulated metabolites across HMDB subsets

### Aging-related biomarker identification

We leveraged the predictive capabilities of MetaKGE to forecast metabolites associated with aging. From the literature, we identified 235 aging-related metabolites. Five metabolites with high identifiers and sparse known information were excluded. The remaining 230 metabolites were divided into a training set of 180 and a test set of 50. The training set included triples of the form {metabolite, related to, Aging}, which were integrated into MetaKGE’s training process. Upon the completion of MetaKGE’s training, we performed predictions for triples of the format {, related to, Aging}.

During this prediction phase, the MetaKGE model assigned a score to each triple consisting of an entity *s* in the form of {s, related to, Aging}. We ranked candidate triples using the trained model’s prediction scores.

We then compared these ranking outcomes with the data in our training and test sets to evaluate the predictive performance of MetaKGE.

## Acknowledgments

This study was funded by the National Natural Science Foundation of China (grant number 6220071694 to J.W. and 32422048 to L.C.), the Non-profit Central Research Institute Fund of the Chinese Academy of Medical Sciences (2021-RC350-008, L.W.), the National Key R&D Program of China (2024YFF0507404 to J.W. and 2022YFA1106300 to L.W.), State Key Laboratory Special Fund (2060204, L.W.)

## Author Contributions

Conceptualization: J.W., L.C., L.W, Y.L. Data acquisition: Y.L., X.G., L.W, L.C., J.W. Analysis: Y.L., X.G., L.W., L.C., J.W. Methodology: Y.L., L.W., L.C., J.W. Supervision: L.W., L.C., J.W. Funding acquisition: L.W., L.C., J.W. Project administration: L.W., L.C., J.W. Writing: Y.L., wrote the manuscript with the help of X.G., L.Q., L.W., L.C., and J.W. All authors discussed the results and reviewed the manuscript.

## Data availability

The MetaKG triples and entities can be downloaded directly from www.metakg.xyz/ download subject to the terms of the underlying data sources. Raw data from HMDB (v5.0) is accessible at https://hmdb.ca/downloads. Raw data from KEGG (release 2023/10), PubChem (records by 2024), and ChEBI (rel217) is obtained via the bioservices package, which can be found at https://github.com/cokelaer/bioservices. The Clintox, BBBP, Delaney, Clearance, and Lipo datasets are hosted at https://huggingface.co/zpn, while the Tox21, PCBA, Freesolv, and HIV datasets are available at https://huggingface.co/HUBioDataLab. The Bitter and Sweet (b&s) dataset can be accessed at https://github.com/cosylabiiit/bittersweet. Pretrained KGE model weights and metabolite embeddings are also available on our http://www.metakg.xyz website.

## Code availability

The MetaKG source code is available under the MIT license on GitHub (https://github.com/YuxingLu613/MetaKG). Jupyter notebooks exemplifying how to use the Meta2Vec embeddings from MetaKG can be downloaded from https://github.com/YuxingLu613/meta2vec/blob/main/example.ipynb.

## AI-assisted editing

OpenAI Codex assisted with language editing, formatting and consistency checks of the existing manuscript. It was not used to generate experimental data or research results.

---

**Extended Figure 1.**
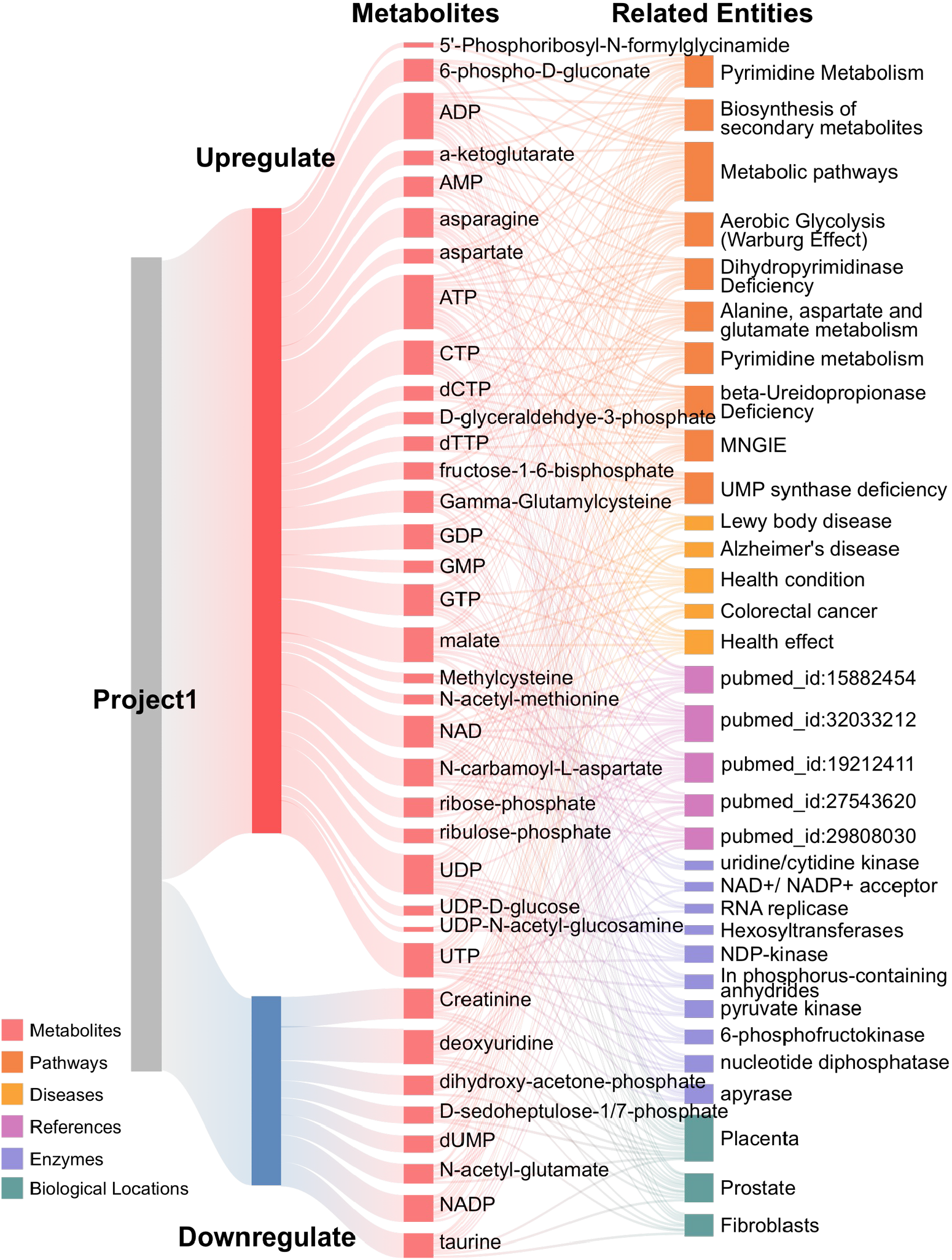
Sankey Plot Analysis Using MetaKG. MetaKG enables visualization statistical analysis and visualizes multi-modal relationships of a set of metabolites using a Sankey plot.

**Extended Figure 2.**
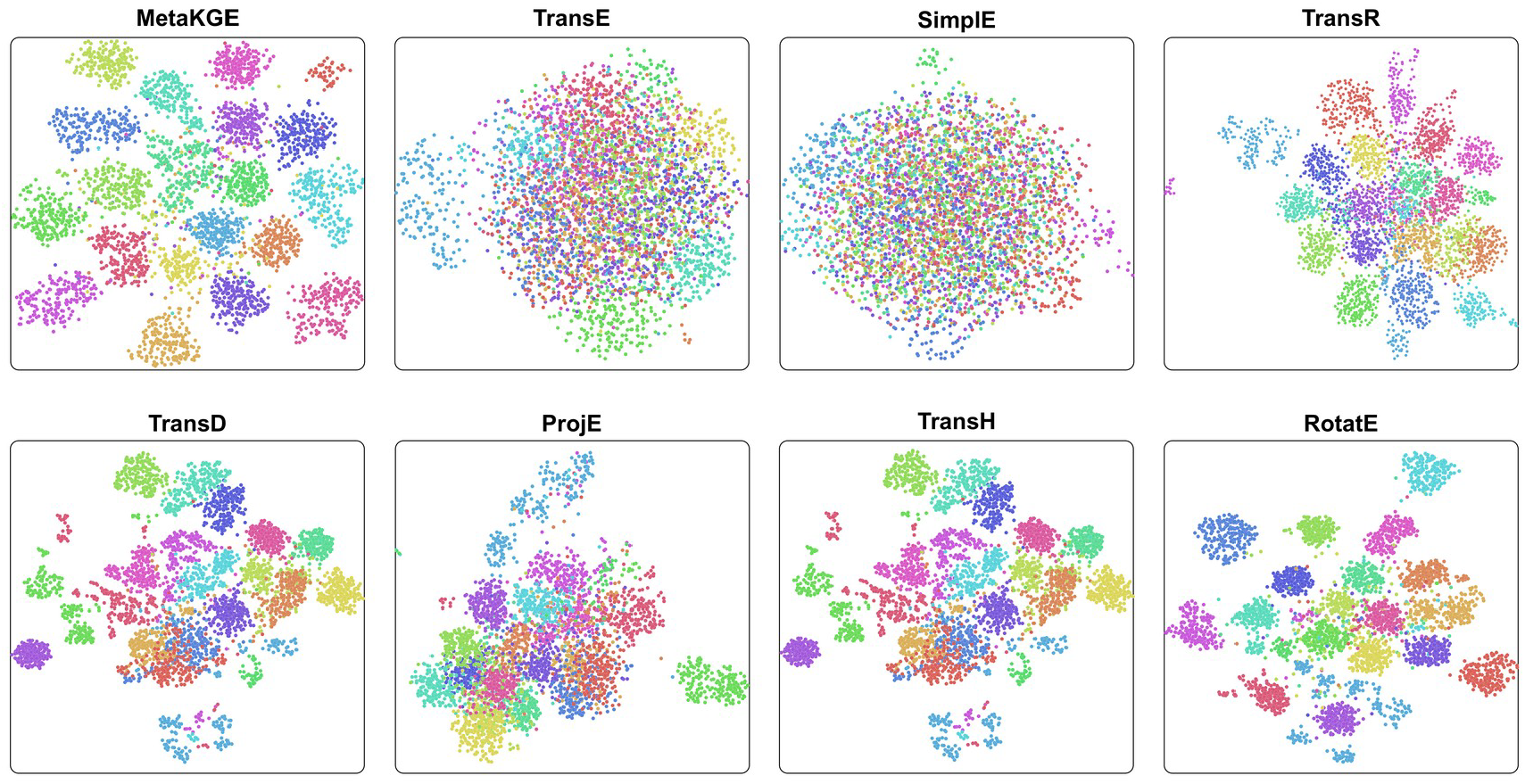
2D Visualization of Different Knowledge Graph Embedding Methods. Through the triple contrastive learning framework, MetaKGE demonstrates a more visually apparent clustering effect compared to other knowledge graph embedding methods, bringing entities of the same category closer together. Additionally, the RotatE method on which MetaKGE is based also achieves a notable clustering effect.

**Extended Figure 3.**
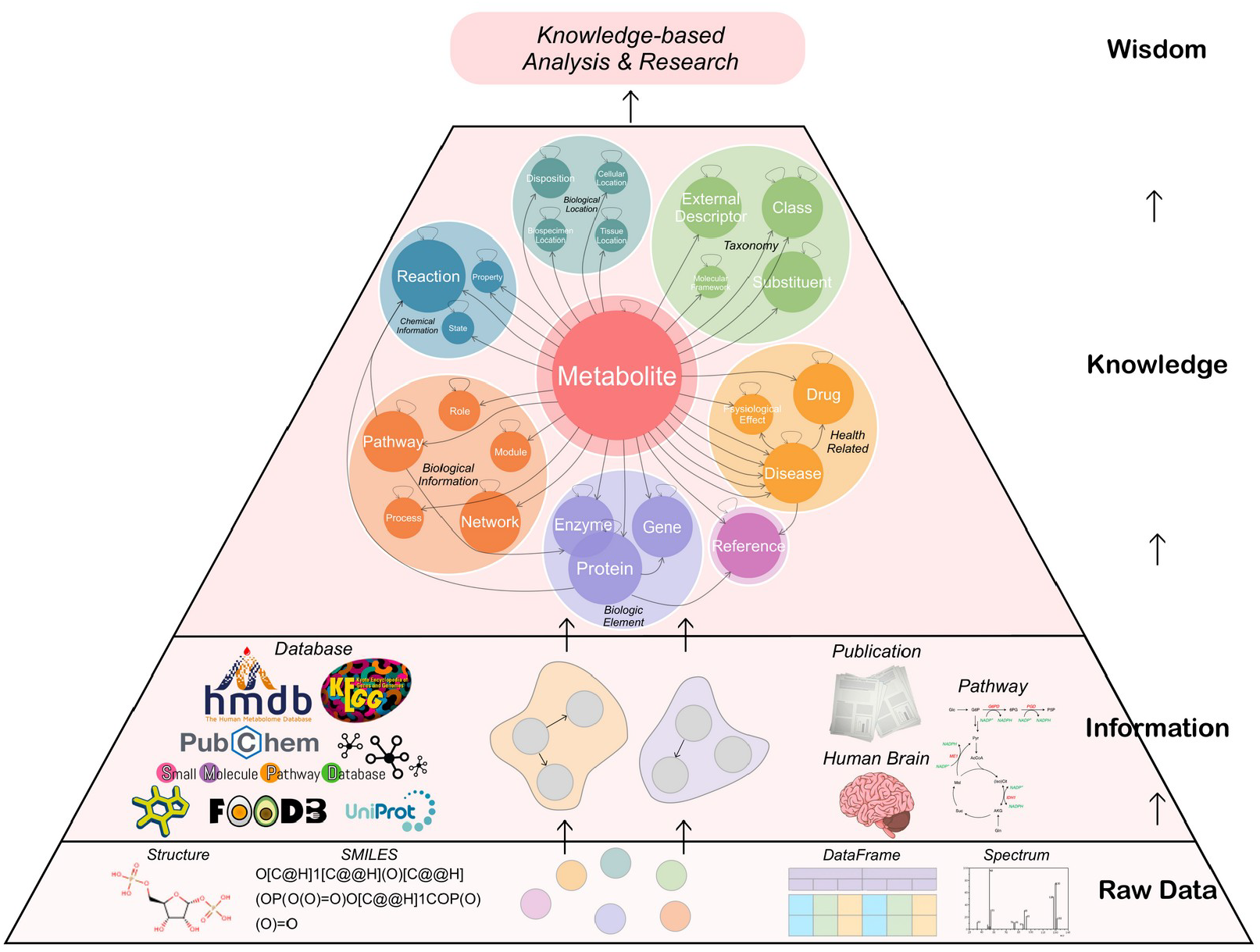
Knowledge Pyramid of MetaKG. The base layer of metabolomics knowledge consists of raw data (including metabolite structure, SMILES sequence, spectrum graph, etc.), and the relationships among these data form biological information (stored in databases and publications). Further integration of these information results in the Metabolite-centric knowledge graph (MetaKG). Harnessing this metabolomics knowledge with AI techniques can produce new wisdoms.

**Extended Figure 4.**
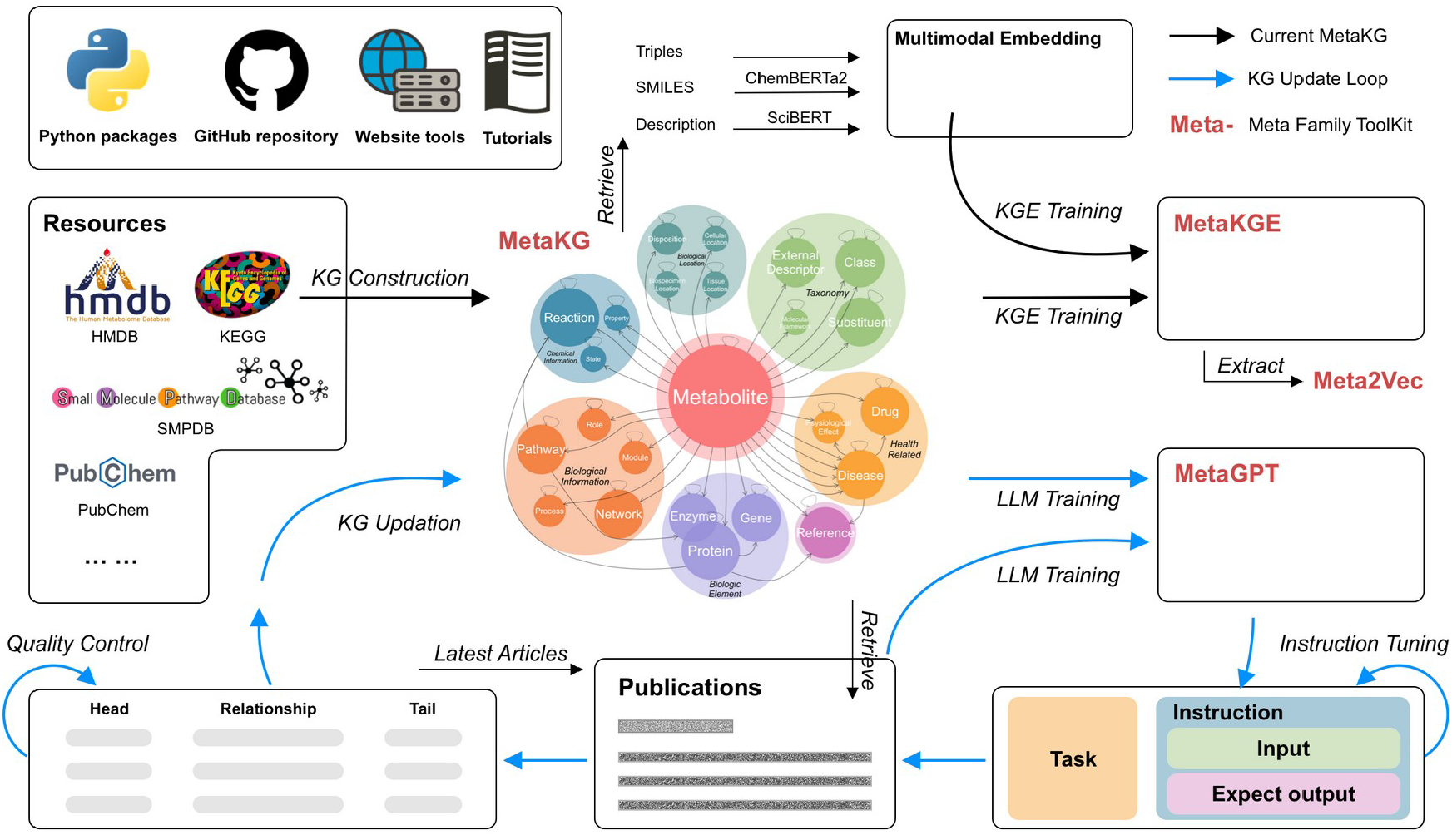
A Prospective on Meta Family Toolkit and MetaKG Updating. We construct MetaKG by integrating several different metabolite databases and obtain a representation (MetaKGE) for each entity, which has proven to be a robust descriptor. Meta2Vec is the extracted metabolite embedding that can be used for downstream prediction of metabolite-related properties. MetaKG-LLM, by retrieving and parsing the latest information using large language models, is integral to the automatic updating of MetaKG.

