## Supplementary Information for "Metabolite-centric Knowledge Graph Unveils Predictive and Mechanistic Insights through Self-supervised Multimodal Learning"

Yuxing Lu *et al.*

Contents: Supplementary Notes 1–6; Supplementary Figures 1–13; legends for Supplementary Tables 1–15. Numerical tables are supplied in the accompanying XLSX workbook.

### Supplementary Notes

#### Supplementary Note 1. Choice of metabolic databases

The selection of databases (DBs) for inclusion in MetaKG was guided by several key criteria. Our primary consideration was database coverage and quality. HMDB (Human Metabolome Database) was selected as the primary reference database due to its comprehensive coverage of human metabolites, detailed biochemical information, and regular updates. HMDB contains >114,000 metabolite entries with extensive metadata including physiological concentrations, chemical classifications, and disease associations. KEGG (Kyoto Encyclopedia of Genes and Genomes) was included for its well-curated metabolic pathway information and standardized compound representations. KEGG's strength lies in its systematic organization of metabolic pathways and reactions, making it invaluable for pathway analysis. SMPDB (Small Molecule Pathway Database) was chosen for its focus on small molecule pathways and disease mechanisms, complementing KEGG's broader pathway coverage with detailed human-specific metabolic pathways.

Database interconnectivity was another crucial factor in our selection. These databases were chosen due to their extensive cross-referencing capabilities: HMDB provides mappings to both KEGG and SMPDB identifiers, KEGG compounds are widely referenced across other biochemical databases, and SMPDB integrates well with both HMDB and KEGG through standardized identifiers.

While other metabolic databases exist, several factors influenced our decision not to include them. MetaCyc, while comprehensive, shows significant overlap with KEGG pathways and has more complex licensing requirements. ChEBI focuses on chemical entities rather than metabolic pathways, while PubChem's scope is too broad, including non-metabolic compounds. LipidMaps, with its specialized focus on lipids only, was too narrow for our purposes.

Technical considerations also played a key role in our selection process. The selected databases offer stable API access, regular updates and maintenance, well-documented data structures, standardized compound identifiers, and machine-readable formats.

Finally, community adoption was a significant factor. HMDB, KEGG, and SMPDB are widely used in the metabolomics community, ensuring broader applicability of MetaKG, easier integration with existing tools, and greater reliability of information due to community curation. This combination of databases provides a balanced coverage of metabolic information while maintaining data quality and interoperability.

**Supplementary Note 2. Data preprocessing**

MetaKG is a comprehensive knowledge graph that integrates metabolomic data from multiple sources, such as the Human Metabolome Database (HMDB), the Kyoto Encyclopedia of Genes and Genomes (KEGG), and the Small Molecule Pathway Database (SMPDB). The construction of this graph necessitated a meticulous and multi-stage data preparation pipeline, encompassing data acquisition, parsing, cleaning, integration, and alignment. The goal of this process was to ensure that all information in MetaKG was harmonized, accurate, and consistent, facilitating meaningful insights into metabolic pathways, disease mechanisms, and therapeutic targets.

**Data Acquisition and Parsing**

The first step in our workflow was data acquisition, where we retrieved metabolomic data from HMDB, KEGG, and SMPDB. These databases store information in a variety of formats, including XML, CSV, and JSON, which required tailored parsers to extract the relevant data efficiently and consistently.

**HMDB Parsing:** The HMDB database provides detailed metabolite data in XML format. First, we converted these XML files into JSON format for easier handling. Using a schema-based parser, we extracted data related to metabolites, their properties, and their associations with diseases, pathways, and biological processes. For instance, the compound Lactate (HMDB0000190) is associated with the metabolic disorder Lactic Acidosis and is participated in Pyruvate metabolism pathway (map00620). This data was parsed into entities and triples as follows:

Entities:

{hmdb_id:HMDB0000190, Metabolite}

{disease:Lactic Acidosis , Disease}

{pathway_id:map00620, Pathway}

Triples:

{hmdb_id:HMDB0000190, has_disease, disease:Lactic Acidosis}

{hmdb_id:HMDB0000190, has_pathway, pathway_id:map00620}

This structure ensures that each relationship is clearly defined, allowing for easy querying and visualization in the graph.

**KEGG Parsing:** KEGG provides metabolomic data through *bioservices* (https://pypi.org/project/bioservices/), which we accessed using programmatic interfaces. These data were categorized into various types, including metabolites, pathways, and enzymes. We utilized *bioservices* to retrieve relevant data and organized it into a standard format. For example, a KEGG entry for the compound Pyruvate (C00022) links it to various metabolic pathways like Tyrosine metabolism (map00350) and the Citrate cycle (map00020), which we captured as:

Entities:

{kegg_id:C00022, Metabolite}

{pathway_id:map00350, Pathway}

{pathway_id:map00020, Pathway}

Triples:

{kegg_id: C00022, has_pathway, pathway_id:map00350}

{kegg_id: C00022, has_pathway, pathway_id:map00020}

These entities and triples were extracted and processed similarly to those from HMDB.

**SMPDB Parsing:** SMPDB provides pathway-related data in CSV format. We employed *pandas* for efficient data manipulation, extracting information about small molecule pathways and their constituent metabolites. Pathway-related entities were mapped to metabolites, and relationships were stored in the standard triple format.

After parsing the data, we stored each extracted element as an entity (name and category) and encoded relationships between these entities as triples. This standardization of entities and relationships provided a foundation for further processing and integration.

**Knowledge Cleaning**

Once the data was extracted, the next step was knowledge cleaning. This stage aimed to refine the data by removing noisy, redundant, or incorrect associations, ensuring that the final knowledge graph only contained high-quality, biologically meaningful relationships.

**Automated Filtering:** We implemented rule-based filters to identify overly broad or generic relationships. For example, associations like "has_effect" with a vague description such as "Health Effect" were flagged and removed. These types of relationships often involve high-degree entities that connect to many other entities, making them non-specific and not informative. By eliminating such relationships, we narrowed down the focus to specific, well-defined associations, like the role of a metabolite in a particular disease or process.

**Manual Curation:** Despite automation, some entities required manual curation due to the complex nature of metabolomic data. For instance, a compound may be associated with several diseases across different databases, but some associations may be irrelevant or outdated. Manual curation by domain experts ensured that each relationship was biologically plausible and relevant.

**Integration and Entity Standardization**

The integration phase focused on reconciling the diverse identifiers and naming conventions across the different data sources. This was critical for ensuring that the knowledge graph was unified and free from ambiguity.

**Cross-Reference Mapping:** We used external cross-reference data from sources like PubChem and UniProt to map entities from different databases to a unified set of identifiers. For example, if the same metabolite like lactic acid was represented by different IDs in HMDB (HMDB0000190) and KEGG (C00186), we created explicit “the_same_as” relationships to link these IDs, thus harmonizing them into a single entity. This allowed us to consolidate knowledge across databases without losing important information.

Example:

{HMDB0000190, the_same_as, C00186}

**Normalization of Entity and Relationship Names:** Variations in formatting, such as different letter casing and use of special characters, were standardized to avoid duplicate entities. We implemented string normalization procedures to ensure that entities with different representations (e.g., "ATP" vs. "atp") were merged into a single standardized form. We also harmonized the various relationship terms used across databases. Different data sources often used different terminology for the same type of relationship. For example, KEGG might use the term "related_to_pathway," while HMDB uses "has_pathway." To ensure consistency, we mapped all pathway-related relationships to a single term, “has_pathway,” across the graph.

**Numerical Data Preprocessing**

In addition to qualitative relationships, MetaKG also integrates quantitative data, such as metabolite concentration levels in various biological contexts. To ensure the reliability of this data, we applied several preprocessing steps. Missing values and outliers were identified using statistical methods such as interquartile range (IQR) filtering. Missing data points were either imputed or discarded depending on the context, while outliers were removed to avoid bias in downstream analyses.

We standardized metabolite concentrations to a common reference scale based on known baseline values in healthy individuals. This standardization was essential for comparing metabolic changes in disease states. For example, if the metabolite Pyruvate (HMDB0000243) shows a concentration of 100 µM in healthy individuals but 300 µM in colorectal cancer patients, the relationship would be encoded as:

{hmdb_id:HMDB0000243, is_upregulated_in, colorectal cancer}

This process allowed us to identify and highlight metabolites that were significantly altered in disease conditions, enabling insights into metabolic disruptions and potential biomarkers.

**Final Knowledge Alignment**

In the final stage, we ensured that all entities and relationships were properly aligned across databases. By mapping metabolites, diseases, and pathways to a unified set of identifiers, we avoided duplication and ensured that each entity was consistently represented. Furthermore, we merged similar or synonymous relationships to reduce redundancy and improve the overall clarity of the knowledge graph.

Example: The relationship "has_disease" was unified across all databases to ensure that all disease associations for a particular metabolite were consistently labeled, regardless of whether the data came from HMDB, KEGG, or SMPDB.

This comprehensive alignment resulted in a clean, integrated dataset that accurately reflected the relationships between metabolites, diseases, pathways, and other biological entities.

**Supplementary Note 3. ChemBERTa-2 and SciBERT models**

**ChemBERTa-2**

ChemBERTa-2 encodes metabolite SMILES sequences using a pretrained chemical language model. MetaKG uses the DeepChem/ChemBERTa-77M-MLM checkpoint and extracts the final [CLS] representation. The 768-dimensional chemical vector is pooled to match the 128-dimensional MetaKG embedding, as described in the main Methods (reference 34).

**SciBERT**

SciBERT encodes metabolite textual descriptions using the allenai/scibert_scivocab_uncased checkpoint. MetaKG extracts the final [CLS] representation and pools the 768-dimensional vector to the embedding dimension. This provides a semantic representation complementary to the chemical representation (reference 35).

The chemical, semantic and relational representations are aligned through triple contrastive learning to generate MetaKGE embeddings.

**Supplementary Note 4. Open world assumption**

Knowledge graphs inherently operate under what's known as the Open World Assumption (OWA), a critical concept that shapes how we interpret and work with metabolite data. Unlike traditional relational databases, where missing information is typically interpreted as false (Closed World Assumption), OWA treats missing information as simply unknown − a distinction that proves particularly valuable when dealing with metabolic pathways and biochemical relationships.

Consider a practical example: if our knowledge graph lacks a connection between metabolite A and enzyme B, we cannot conclude this interaction doesn't exist; we merely acknowledge that we haven't yet documented this relationship. This perspective is especially relevant in metabolomics, where new interactions and pathways are continuously discovered, and our understanding of metabolic networks remains incomplete.

The implications of OWA become particularly apparent when querying metabolite-enzyme interactions. For instance, when asking "Which enzymes catalyze the conversion of glucose-6-phosphate?", the returned results represent our current knowledge rather than a definitive, complete set. This nuance is crucial for researchers interpreting query results and designing experiments, as it encourages exploration of potentially unknown interactions.

In the context of metabolic pathway analysis, OWA allows for the natural evolution of our knowledge base. As new metabolites are identified and novel enzymatic reactions are characterized, we can seamlessly integrate these discoveries without contradicting existing information. This flexibility is essential given the dynamic nature of metabolomics research and the continuous refinement of our understanding of cellular metabolism.

However, working under OWA also presents certain challenges. Validation of metabolic pathways requires explicit constraints rather than simple absence checks. For example, to assert that a particular metabolite cannot participate in a specific reaction, we must explicitly state this constraint rather than rely on the absence of a relationship. This approach, while more complex, better reflects the reality of biological systems and our evolving understanding of metabolic networks.

### Supplementary Note 5. FAIR principles and resource features

MetaKG is designed as an open resource designed to support FAIR principles to enhance metabolomics research through accessibility, interactivity, and predictive capabilities. This note outlines its implementation of FAIR principles (Findable, Accessible, Interoperable, Reusable) and describes its practical implementation.

**Findable:** MetaKG is hosted at http://www.metakg.xyz, a publicly searchable domain. Each entity and relationship are indexed and searchable via the website's query interface. A machine-readable schema **(Supplementary Figure 1)** ensures discoverability by search engines and repositories.

Accessible: MetaKG source code, tutorials, reproducibility reports and data-access instructions are provided at https://github.com/YuxingLu613/MetaKG. The source code is distributed under the MIT license. Reused source data remain subject to the terms of their respective providers. The demonstration website is described in Supplementary Note 6.

**Interoperable:** MetaKG uses standardized identifiers (HMDB IDs) and ontologies (e.g., disease, taxonomy) for entities, enabling integration with external tools. Relationships are encoded as triples and embeddings are provided in *NumPy* format for computational interoperability. The website uses *echarts* to better visualize the relationship between metabolites **(Supplementary Figure 6)**.

**Reusable:** Data provenance is tracked with curation steps and training details **(Supplementary Figure 7)** fully documented. Users can extend MetaKG with custom datasets (e.g., 235 aging metabolites; **Figure 5b, Supplementary Table 14)**, retrain MetaKGE, and share contributions via GitHub, fostering community reuse. Licensing and metadata ensure legal and scientific reusability.

**Interactive and Predictive Features:** The website offers real-time search (e.g., lactic acid pathways), subgraph visualization, and enrichment analysis. Predictive tools leverage MetaKGE embeddings for tasks like pathway ranking and biomarker identification. These features empower researchers to explore hypotheses interactively, as demonstrated in case studies.

MetaKG's use of FAIR principles and user-centric design make it a dynamic platform, transforming static metabolomics data into actionable, hypothesis-driven insights.

### Supplementary Note 6. MetaKG website introduction

The MetaKG website (https://www.metakg.xyz) offers an intuitive, interactive platform for exploring and analyzing its metabolite-centric knowledge graph, empowering researchers with tools for search, analysis, prediction, path inference, and visualization. Below, we detail its core functionalities, designed for accessibility and scientific discovery.

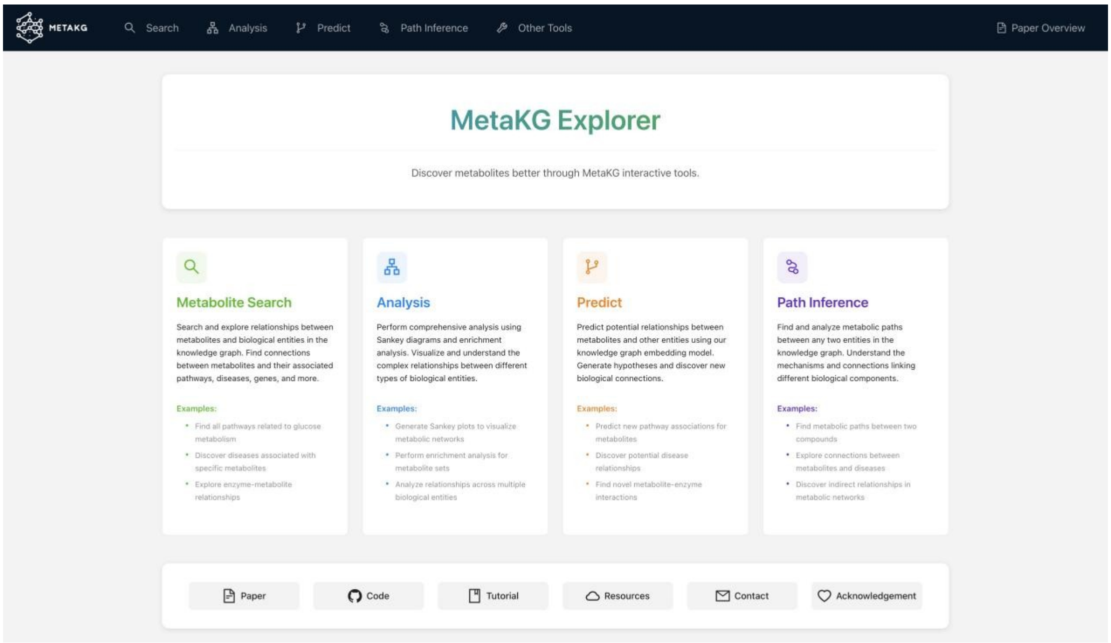

**Search Module:** The search interface enables users to query metabolites and explore their relationships with biological entities (e.g., pathways, diseases). An auto-complete feature simplifies entity selection (e.g., "HMDB0000122#d-glucose"), and results display associated entities. Users can filter by entity type and relationship (e.g., has_disease, has_pathway), facilitating targeted exploration.

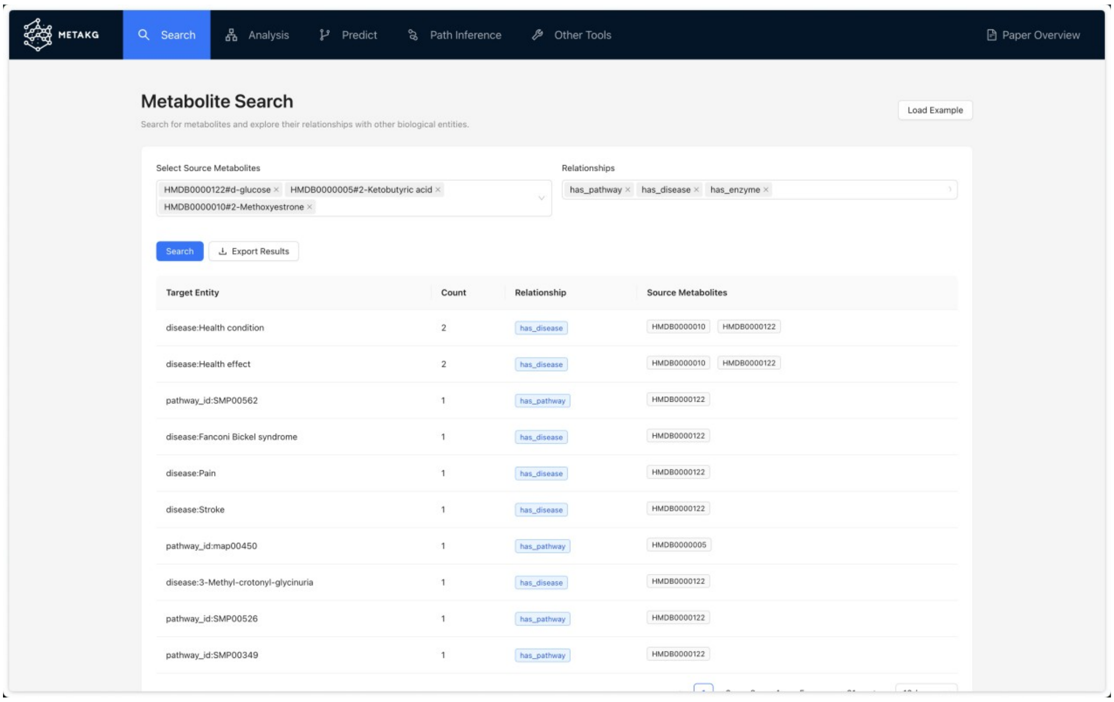

**Analysis Module:** This module provides tools for dissecting metabolic relationships. The Multimodal Sankey Plot visualizes entity connections (e.g., metabolites to pathways like SMPDB00535), supporting multiple types of entities. The enrichment analysis tool performs statistical analysis across multiple modalities, supporting custom entity lists and generating interactive visualizations of enrichment results. This feature is essential for identifying significant patterns and associations within metabolic datasets.

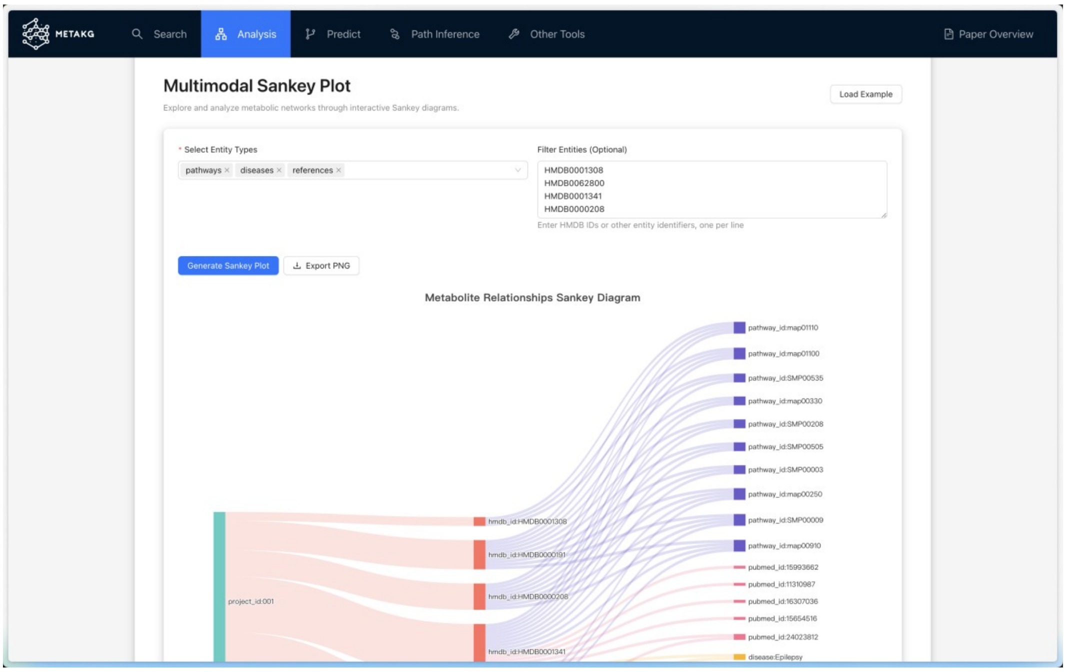


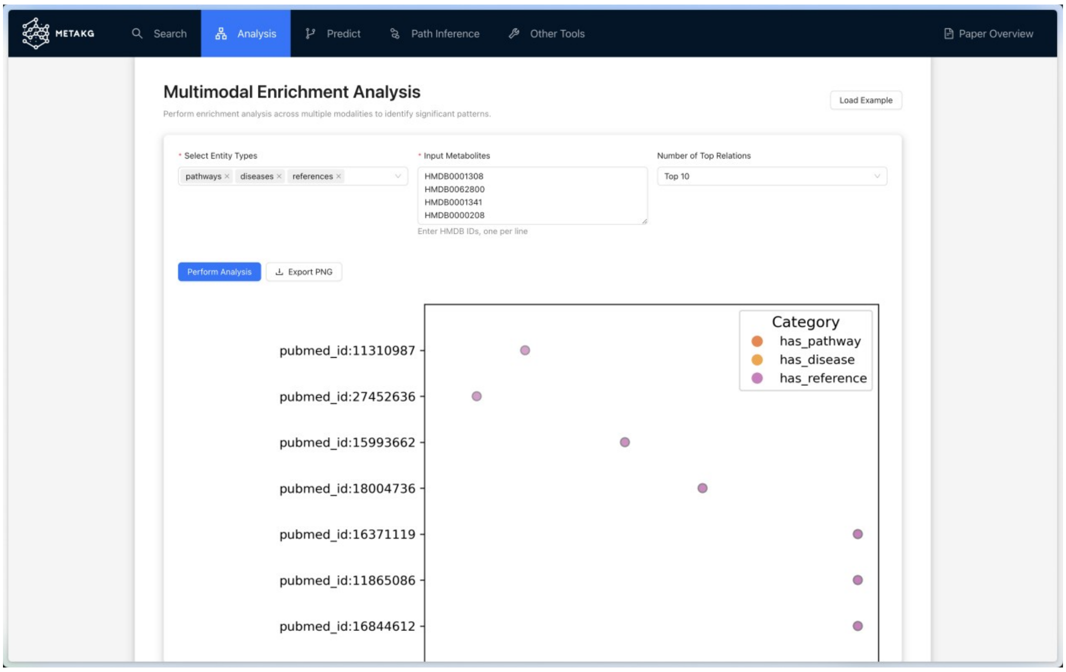

**Prediction Module:** Leveraging MetaKGE embeddings, this module predicts the missing part of a triple. Users can specify head entity, relationship (e.g., has_disease), or tail entity, with confidence scores provided (e.g., 95.2% for Alzheimer's disease).

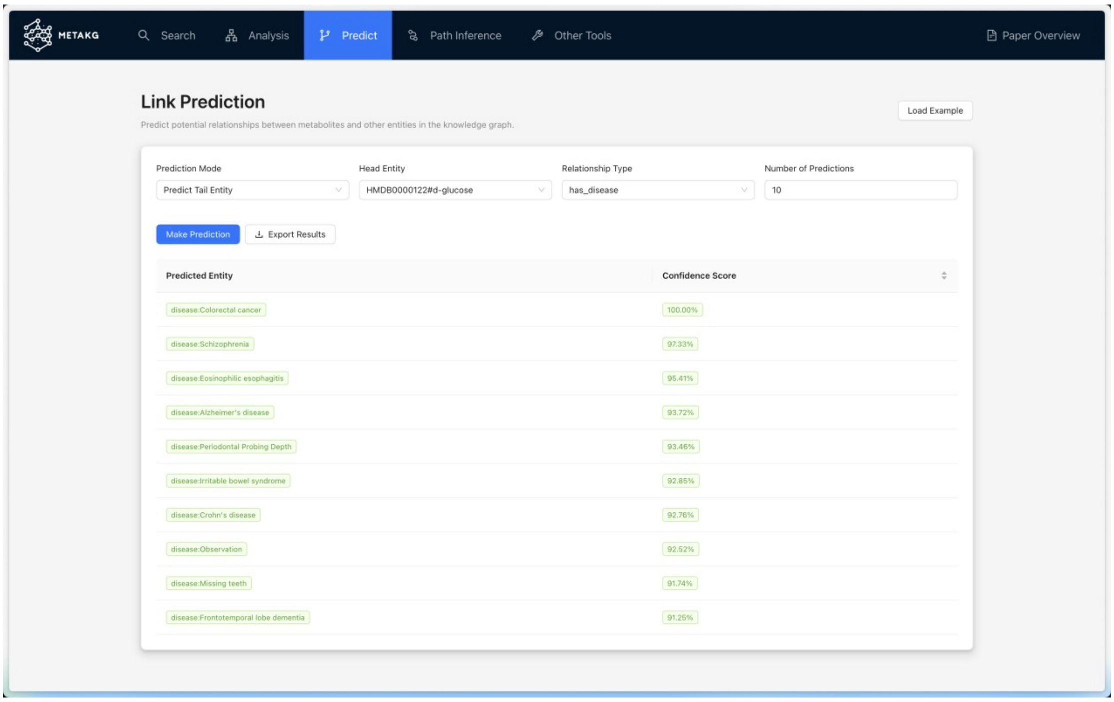

**Path Inference Module:** This tool identifies metabolic paths between entities, with adjustable path lengths and number of paths. For example, paths from NADPH (HMDB0000221#nadph) to dUMP (HMDB0001409#dump) reveal intermediates like EC 3.1.3.- and reaction R02102 (score: 0.7365). Scores rank paths by relevance, aiding in mechanistic discovery.

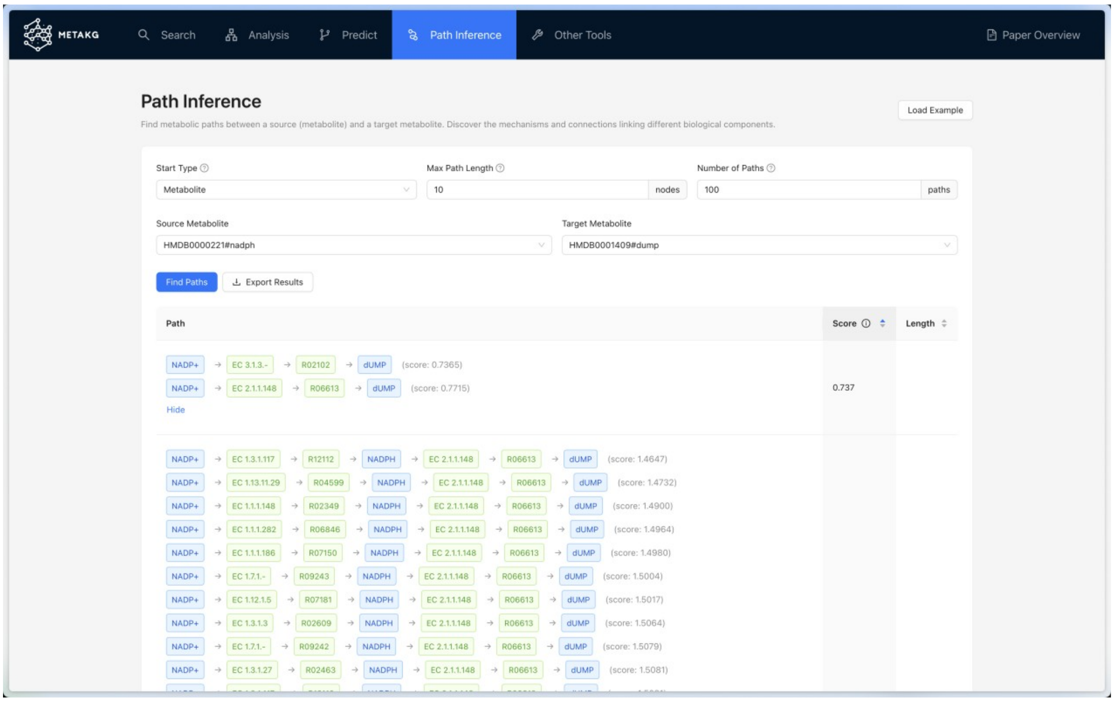

**Visualization tools:** MetaKG provides interactive visualization tools for network analysis and data presentation, with customizable node and edge displays, supporting large-scale networks while maintaining performance. All visualizations can be exported in various formats for publication or further analysis.

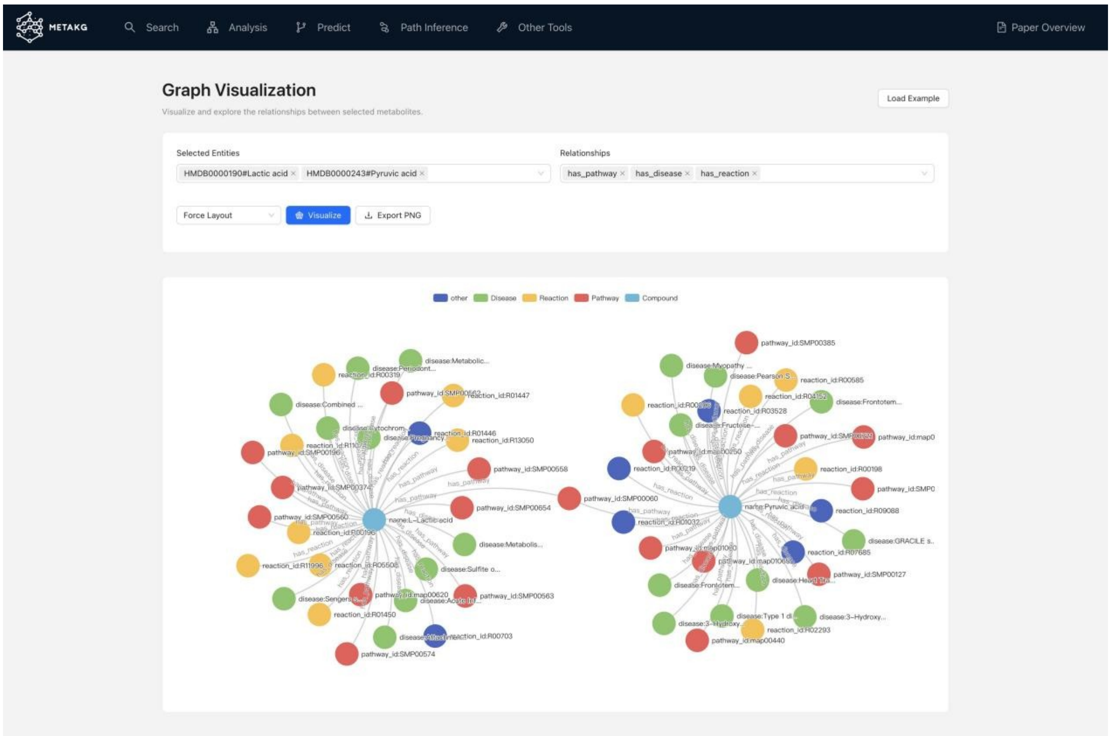


In summary, the MetaKG website integrates these features into a user-friendly interface, supporting real-time exploration and predictive insights, as demonstrated in aging biomarker predictions.

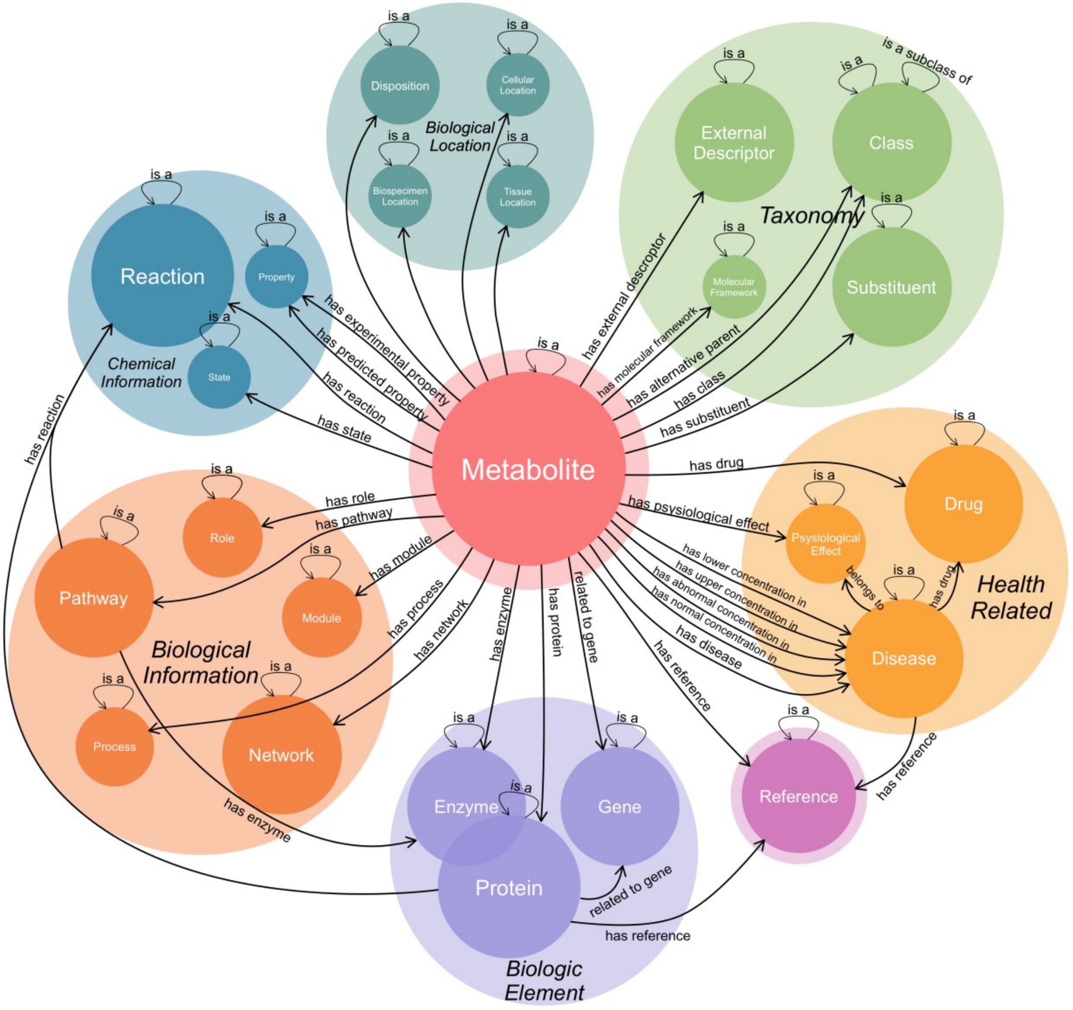


**Supplementary Figure 1. The overall schema of MetaKG.** The metabolite entity is placed at the center of MetaKG, with the remaining entities categorized into seven classes. Entities are self-connected or inter-connected with different relationships.

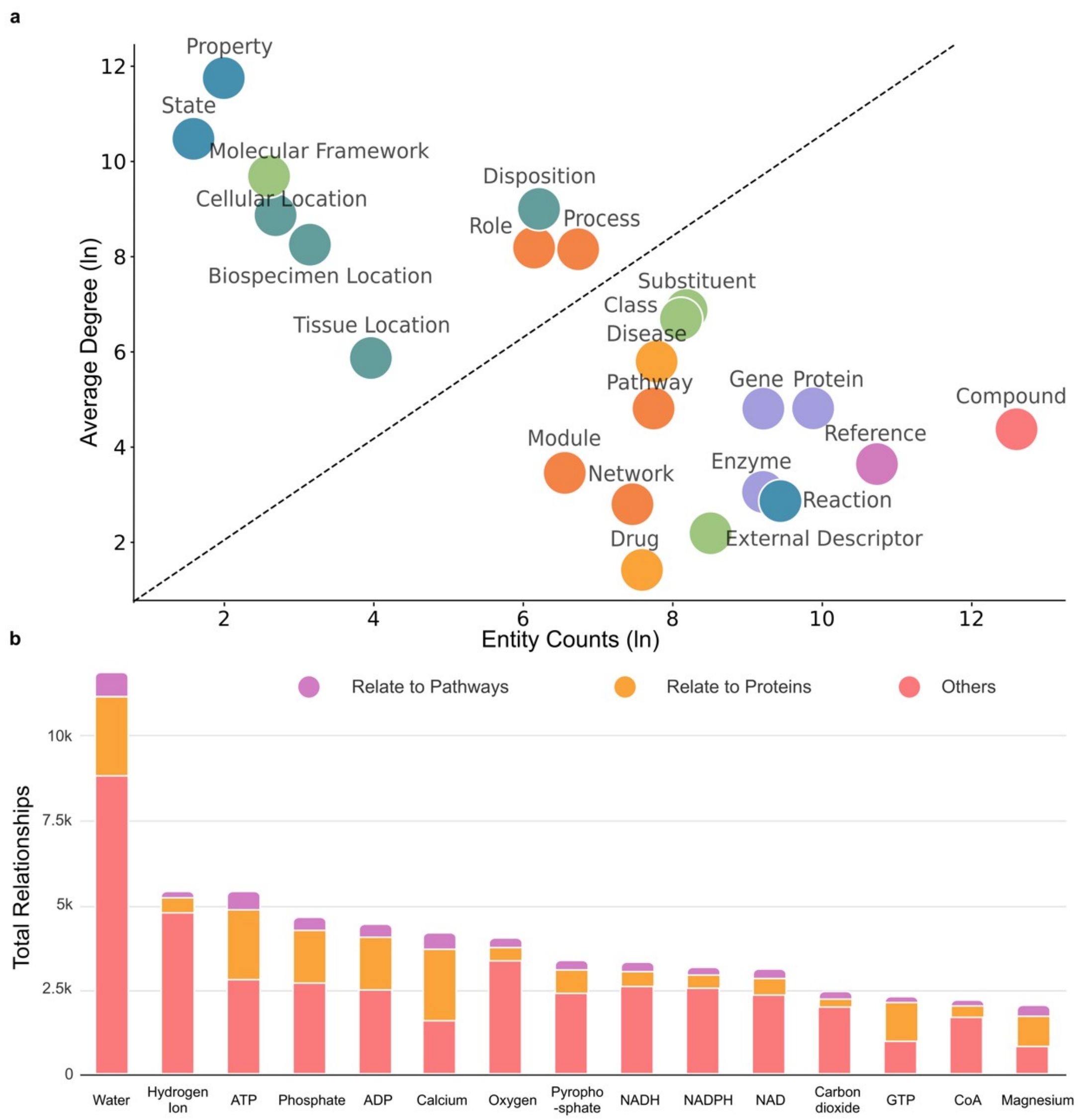


**Supplementary Figure 2. MetaKG statistics. a)** The horizontal axis indicates the logarithmic count (base 10) of entities contained within each category. The vertical axis shows the logarithmic average degree (base 10) of all entities in each category. **b)** Top 15 most frequent metabolites in MetaKG. The most common metabolites are water, hydrogen ions, ATP, and others that are widely involved in various biochemical reactions.

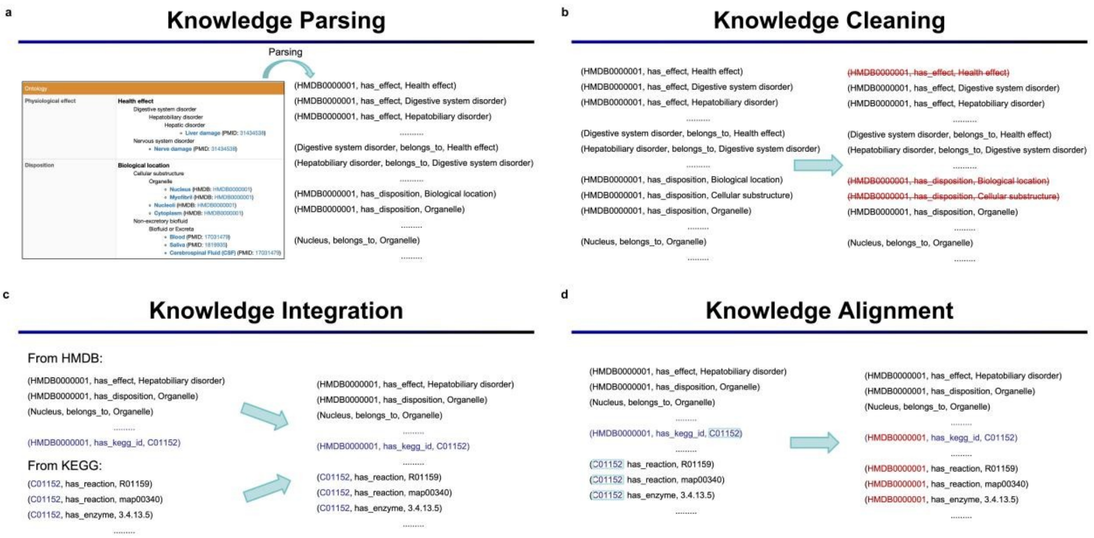


**Supplementary Figure 3. Preprocessing of metabolite knowledge.** **a)** The information stored in different metabolomic databases are parsed into triples. This structured format facilitates the transformation of unstructured biomedical knowledge into a machine-readable format that can be used for further MetaKG construction. **b)** Redundant or incorrect associations are identified and removed. The cleaned data retains only the accurate and relevant associations between entities. **c)** Due to the presence of external link information for a certain entity in the database, the same entities across different databases can be integrated based on their respective identifiers, thereby achieving integration at the database level. **d)** Different descriptions of the same entity in different databases need to be integrated. In MetaKG, we use HMDB id as the unique id for each metabolite.

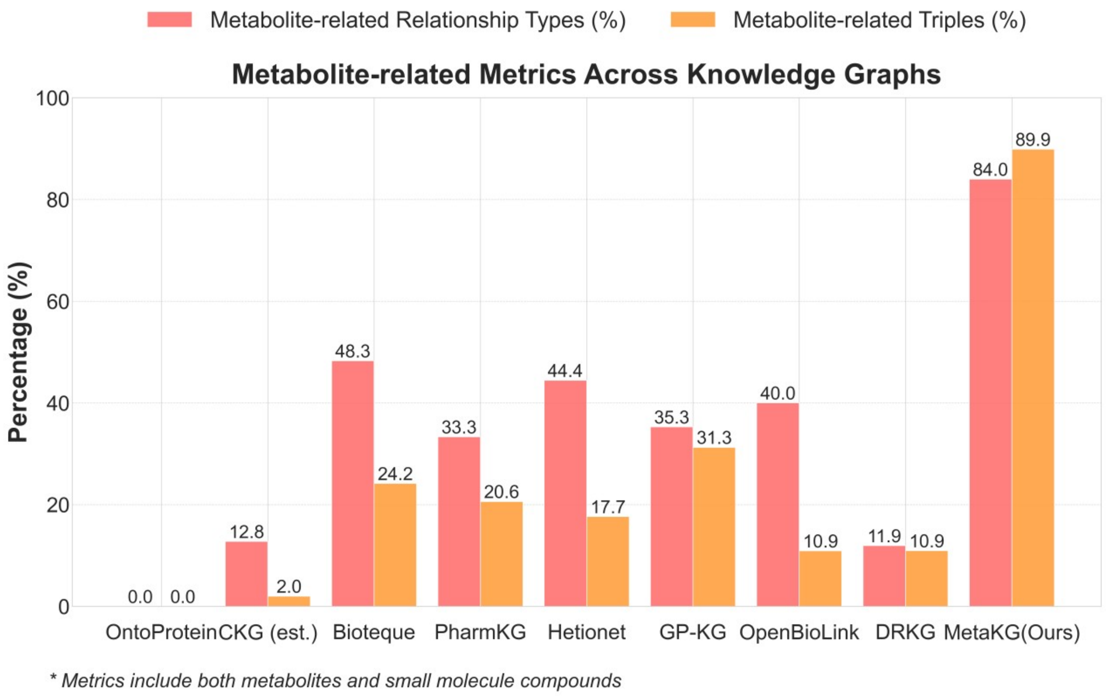


**Supplementary Figure 4. Comparison with other biomedical KGs.** MetaKG has the most metabolite-related relationships and the most metabolite-related triples.

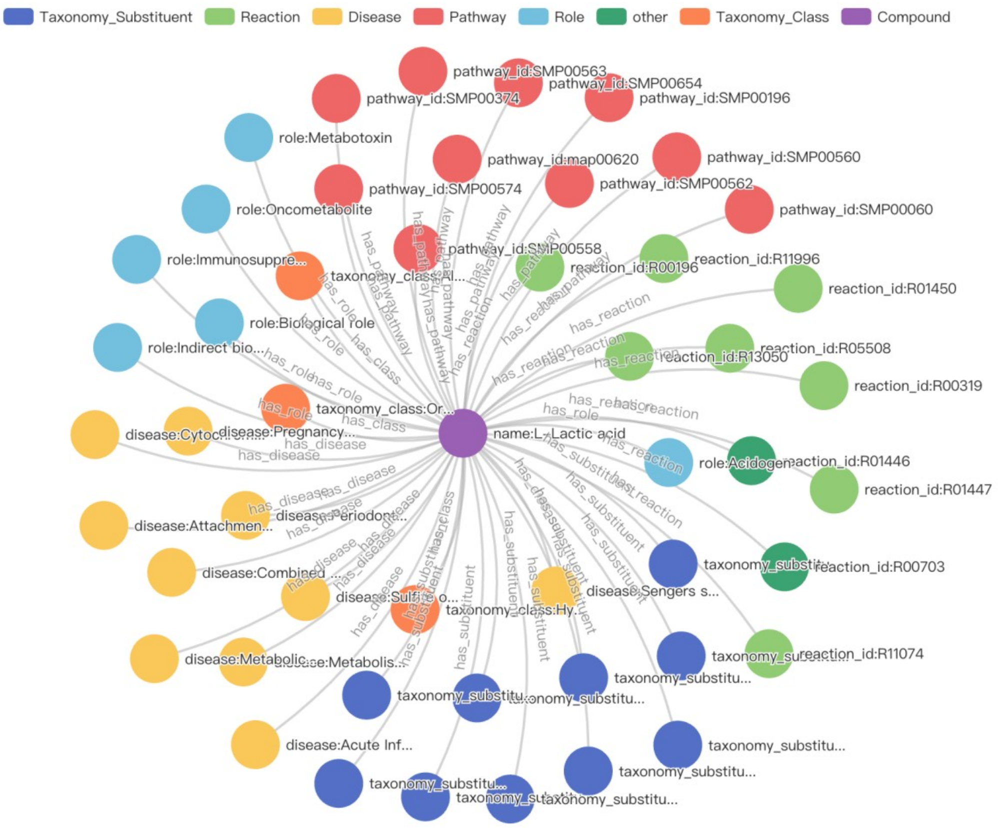


**Supplementary Figure 5. Visualization of Lactic Acid using MetaKG.** We search for all metabolite knowledge related to Lactic Acid using MetaKG and visualize the results.

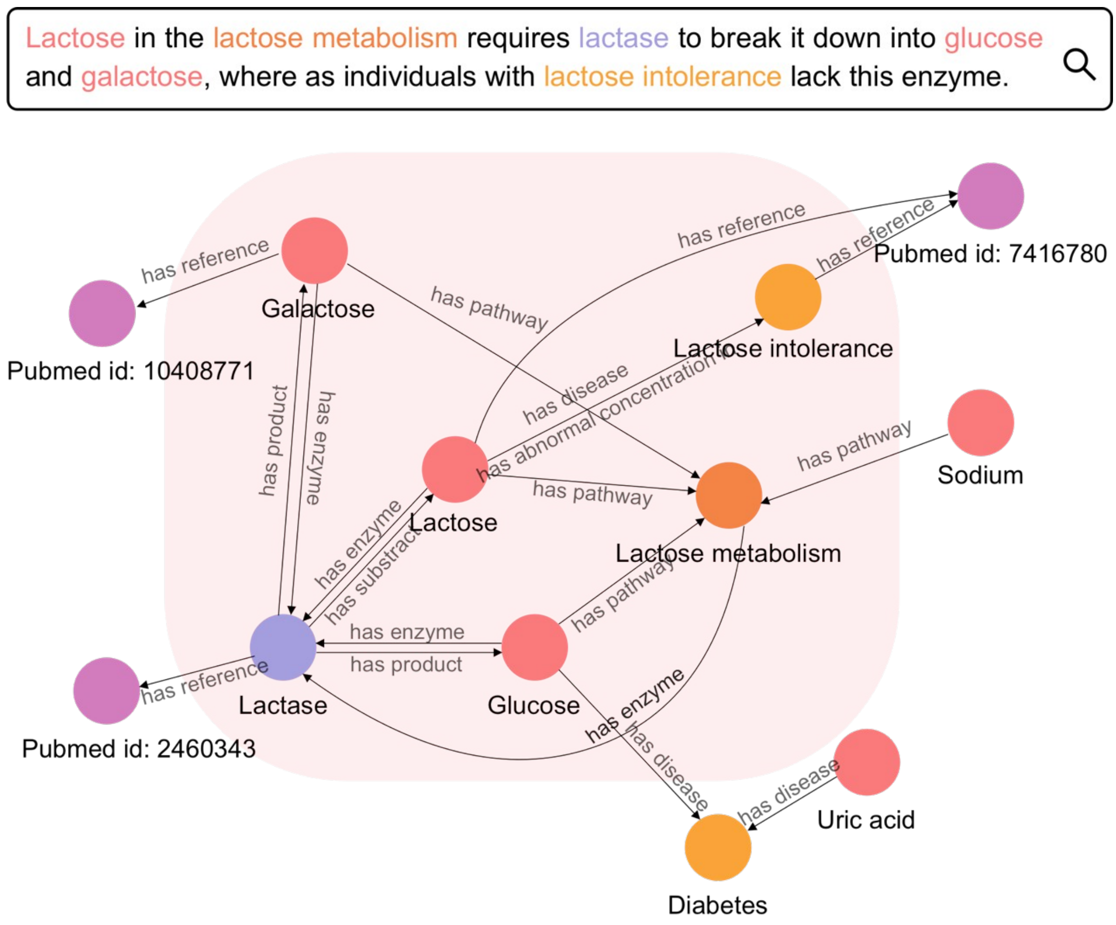


**Supplementary Figure 6. MetaKG knowledge parsing and identification.** MetaKG can parse text to map internal associations and extend the knowledge.

**Algorithm 1** MetaKGE

**Require:** Knowledge Graph $\mathcal{G}$, Embedding Dimension $d$, Learning Rate lr, Number of Multimodal Training Epochs epochs1, Number of Combined Training Epochs epochs2

**Ensure:** Entity Embeddings $E$, Relation Embeddings $R$

$E \leftarrow$ Randomly initialize matrix of size (num_entities $\times d$)
$R \leftarrow$ Randomly initialize matrix of size (num_relations $\times d$)
▷ Multimodal Training Phase
**for** epoch = 1 to epochs1 **do**
  **for** each batch in $G$ **do** ▷ Initialize TCL loss
    $L \leftarrow 0$ ▷ Initialize batch loss
    **for** each triplet $(h, r, t)$ in batch **do**
      $h_{emb}, r_{emb}, t_{emb} \leftarrow$ Retrieve from $E$ and $R$
      $h_{emb\_rot} \leftarrow h_{emb} \cdot r_{emb}$ ▷ Rotate $h_{emb}$ by $r_{emb}$
      $dist \leftarrow h_{emb\_rot} - t_{emb}$ ▷ Compute distance
      $score \leftarrow ||dist||_1$ ▷ Compute L1 norm
      $L \leftarrow L + score$ ▷ Update batch loss
    **end for**
    Perform gradient descent on $L$ to update $E$ and $R$
    $L_{TCL} \leftarrow 0$
    **for** each $m_{id}$ in all_metabolites **do**
      $m_{emb}, m_s, m_t \leftarrow$ Compute using $m_{id}$
      $L_{TCL} \leftarrow L_{TCL} + TripleContrastiveLoss(m_{emb}, m_s, m_t)$
      $m_{emb} \leftarrow (m_{emb} + m_s + m_t)/3$ ▷ Update $m_{emb}$
    **end for**
    Perform gradient descent on $L_{TCL}$ to update $E$ and $R$
  **end for**
**end for**
▷ Combined Training Phase
**for** epoch = 1 to epochs2 **do**
  **for** each batch in $G$ **do**
    $L \leftarrow 0$ ▷ Initialize batch loss
    **for** each triplet $(h, r, t)$ in batch **do**
      $h_{emb}, r_{emb}, t_{emb} \leftarrow$ Retrieve from $E$ and $R$
      $h_{emb\_rot} \leftarrow t_{emb} \cdot r_{emb}$ ▷ Rotate $s_{emb}$ by $p_{emb}$
      $dist \leftarrow h_{emb\_rot} - t_{emb}$ ▷ Compute distance
      $score \leftarrow ||dist||_1$ ▷ Compute L1 norm
      $L \leftarrow L + score$ ▷ Update batch loss
    **end for**
    Perform gradient descent on $L$ to update $E$ and $R$
  **end for**
**end for**
**return** $E, R$ ▷ Output the final embeddings

**Supplementary Figure 7. The overall MetaKGE training pseudo-code.** In general, the MetaKGE training algorithm consists the following modules: multimodal representation acquisition, triple contrastive learning, and dual-phase training. First, the multimodal representation acquisition module utilizes two distinct large language models to produce two different representations of metabolites. Then, the triple contrastive learning models aligns different representations of the same metabolite into a fixed-dimension vector space while maintaining the richness of information from each modality. Finally, the dual-phase training approach guarantees a holistic and in-depth learning experience by concentrating on refining the representations.

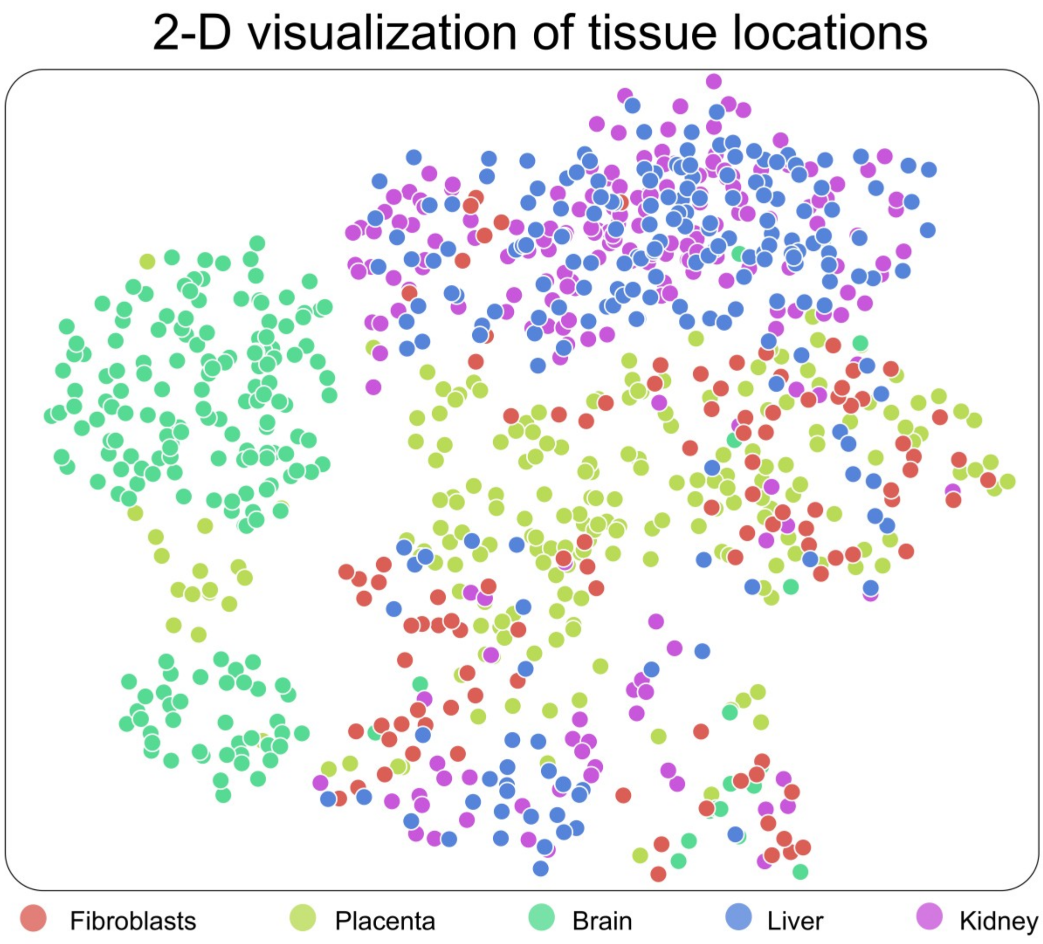


**Supplementary Figure 8. 2-D visualization of metabolites of different tissue locations.** Each point in the visualization likely represents a metabolite, and the color coding indicates the tissue of origin of each metabolite. The proximity of points to each other can suggest similarities in their properties or concentrations between the tissues.

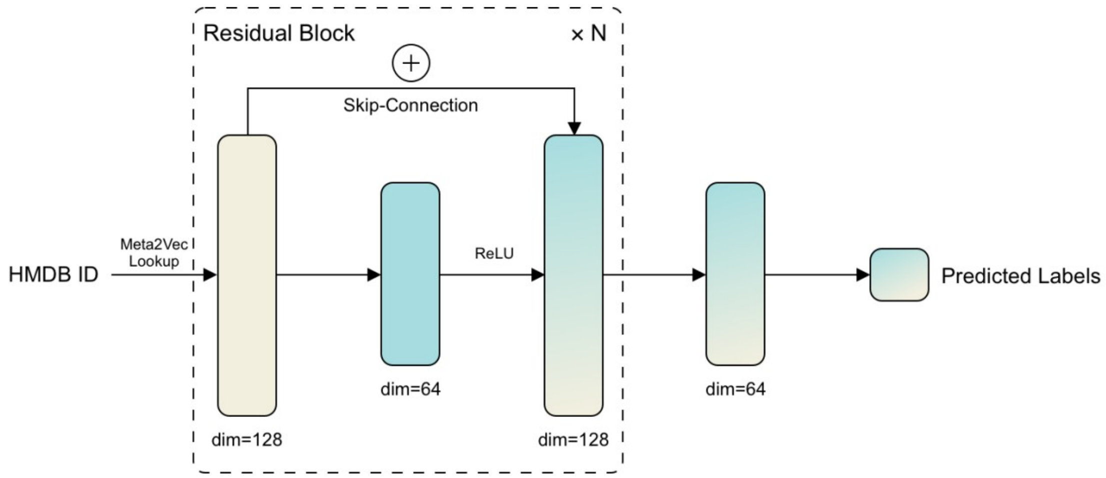


**Supplementary Figure 9. Metabolite external validation model.** We use a deep neural network designed based on the skip-connect architecture to perform training and validation of MetaKGE for assessing the effectiveness of MetaKGE in capturing various characteristics of metabolites.

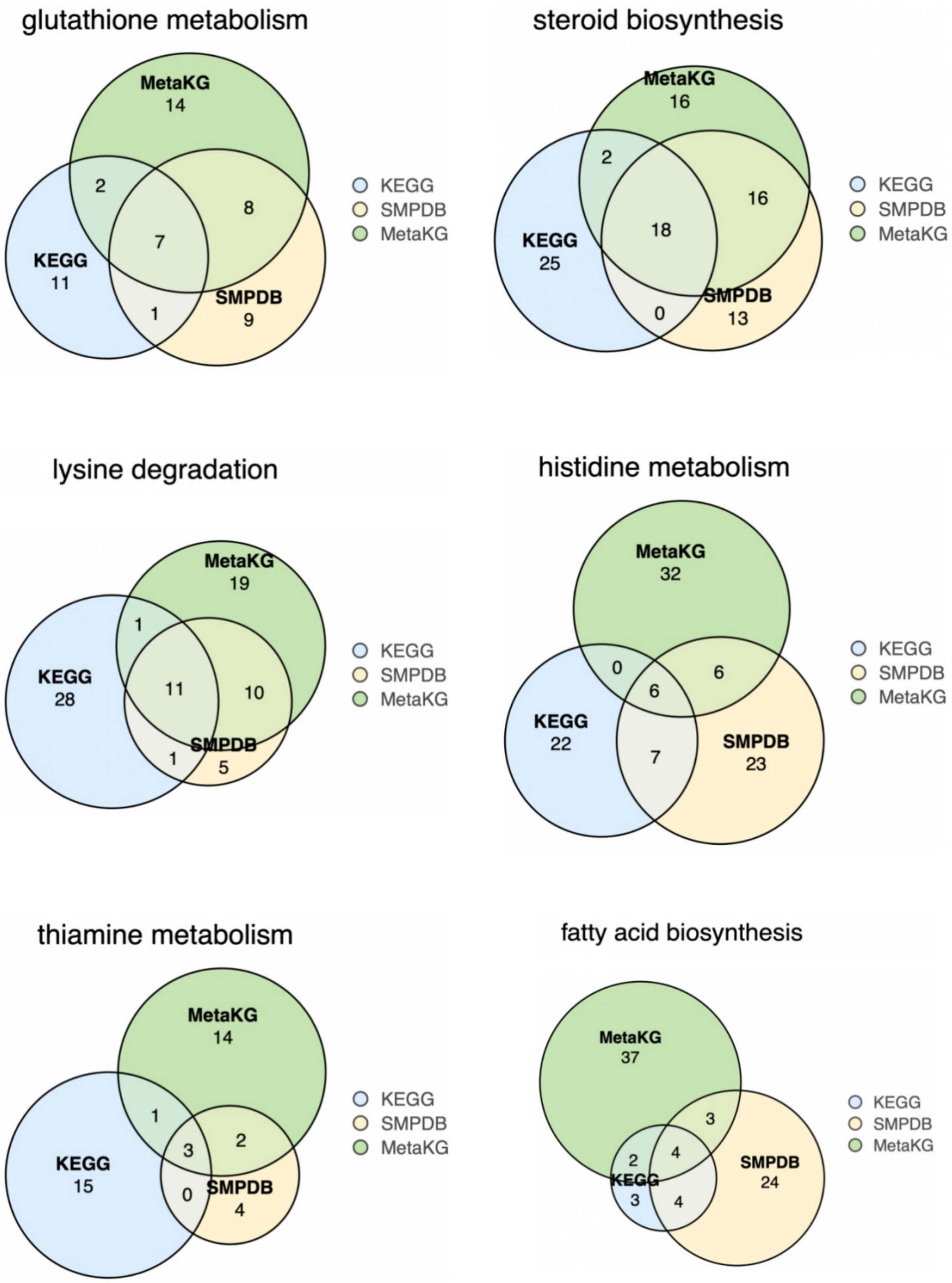


**Supplementary Figure 10. Venn diagrams depicting the metabolite set predicted by MetaKGE with pathways recorded in KEGG and SMPDB.**

**a**

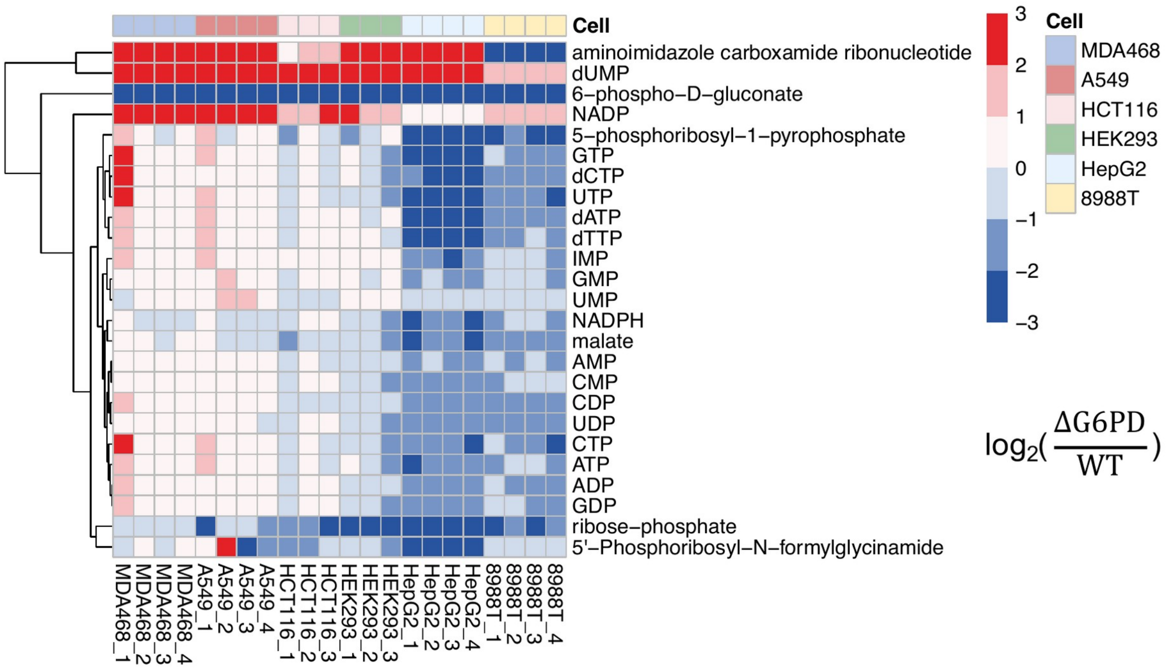


**b**

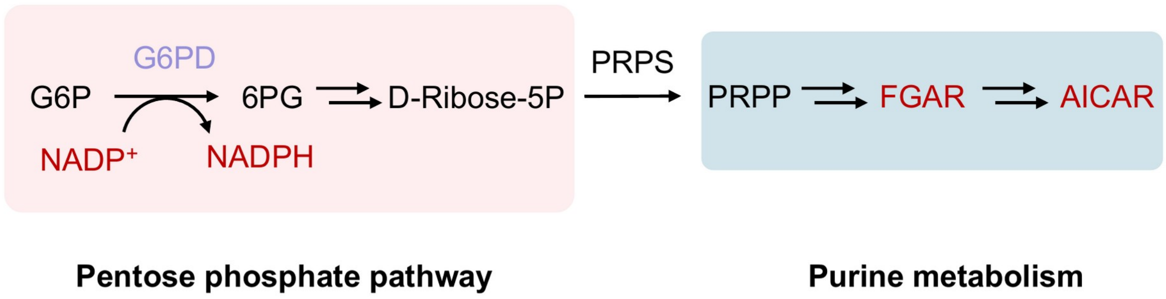


**Supplementary Figure 11. Metabolic alterations in G6PD knockout cells and the classical perturbations to purine metabolism pathways induced by G6PD KO. a) Heatmap of published data reanalyzed from Chen et al. (2019; main-text reference 41), showing** the levels of metabolites (normalized to respective WT cells) in different cell lines with G6PD knockout. **b)** The OxPPP and the de novo purine biosynthesis pathway. Ribose-5-phosphate (Ribose-5P) is the downstream metabolite of oxPPP. Phosphoribosyl pyrophosphate (PRPP) is the beginning metabolite for purine de novo biosynthesis pathway. Phosphoribosyl pyrophosphate synthetase (PRPS) catalyzes conversion of Ribose-5P to PRPP. 5′-Phosphoribosyl-N-formylglycinamide (FGAR) and aminoimidazole carboxamide ribonucleotide (AICAR) are intermediate metabolites in the de novo purine biosynthesis pathway.

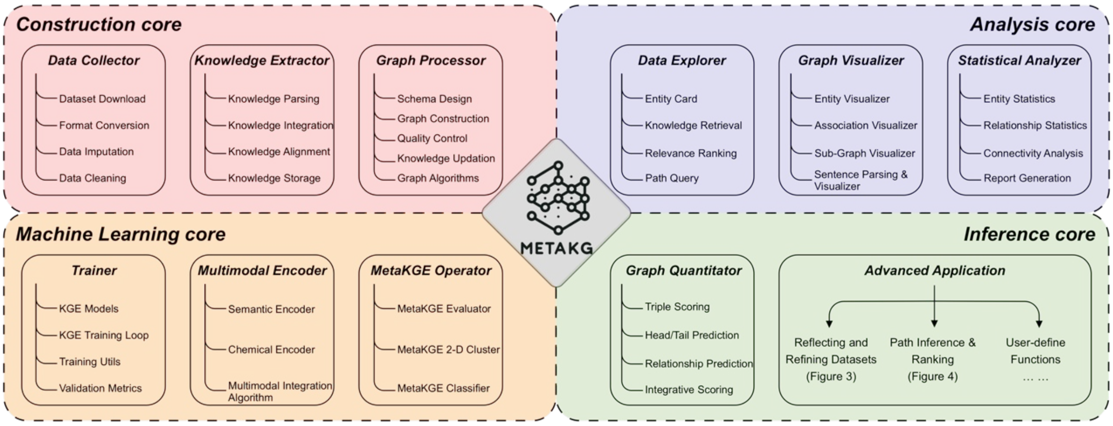


**Supplementary Figure 12. MetaKG repository.** MetaKG is built around four key components: the construction core, analysis core, machine learning core, and inference core.

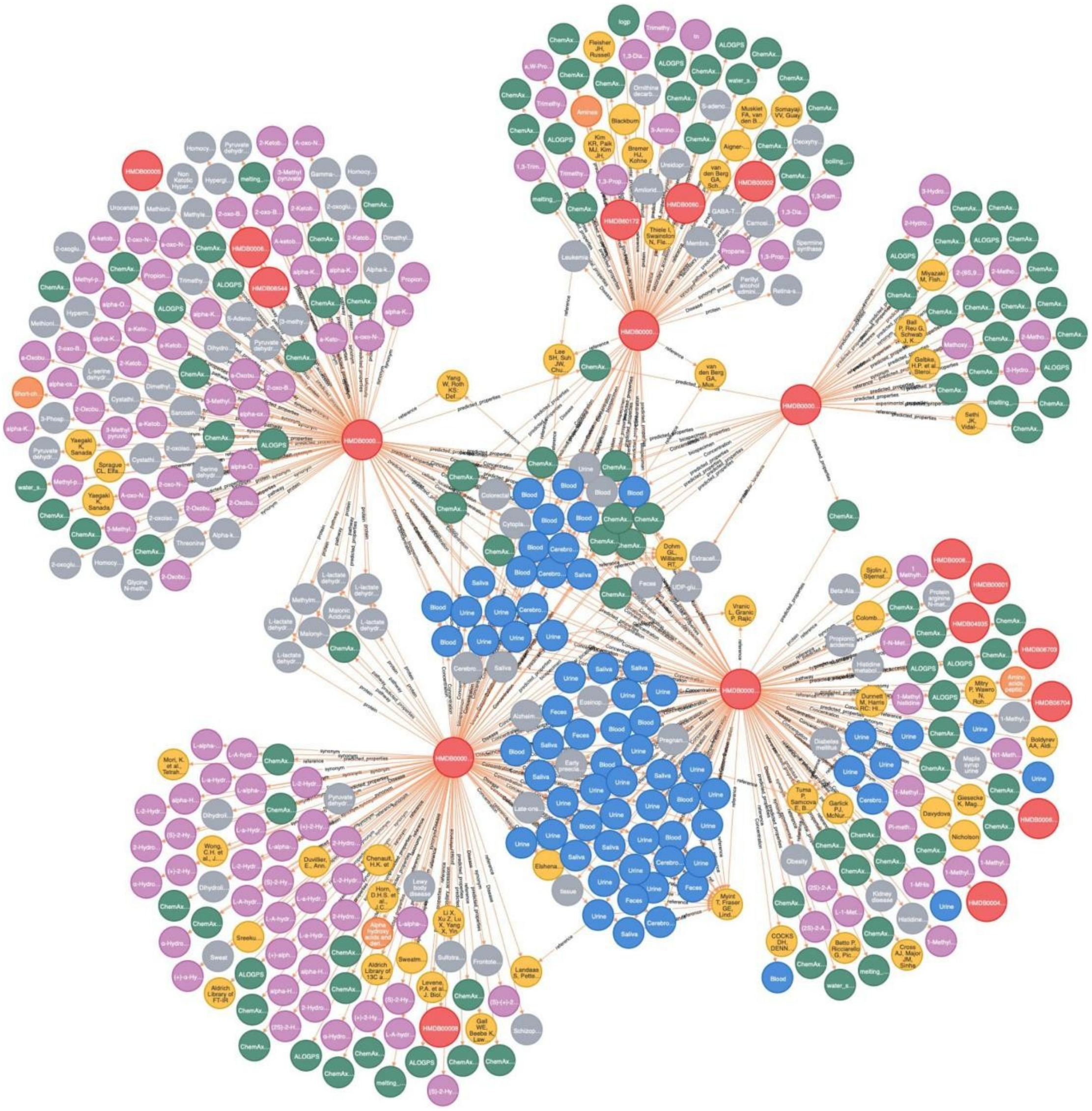

**Supplementary Figure 13. Neo4j as a tool to visualize MetaKG.** Users can use Neo4j as a desktop visualization tool and a graph database manager to provide comprehensive solution for navigating and administrating the MetaKG dataset.

### Supplementary Tables

**Supplementary Table 1. Different entity types and their origin databases.**

**Supplementary Table 2. Different relationship types and their origin databases.**

**Supplementary Table 3. Comparison of different knowledge graph embedding foundation models in MetaKGE.** We compare several different foundational KGE methods and selected the best performing knowledge graph embedding methods and applied it as the base model for our multimodal contrastive learning algorithm. Results are presented in terms of Hit@n, mean rank (MR), MRR (Mean Reciprocal Rank), and clustering coefficient.

**Supplementary Table 4. Different external validation databases and their URLs.**

**Supplementary Table 5. A unified list of metabolites exhibiting high MetaKG relevance scores.** The metabolite sets for pathways with identical names in KEGG and SMPDB as represented in MetaKG, KEGG, and SMPDB, along with the corresponding MetaKG relevance scores for these metabolites.

**Supplementary Table 6. List of metabolites exhibiting potential taxonomic improvement.** The metabolites (from the top 10,000 HMDB IDs) with a difference between the top-1 predicted taxonomy score and the recorded taxonomy score exceeding our provided example are displayed, with the top 10 metabolites with the highest differences highlighted for each taxonomy level.

**Supplementary Table 7. KEGG and SMPDB pathway enrichment results of G6PD KO metabolomics data.** Folate-related pathways rank lower, ranking 39th in KEGG and 36th in SMPDB.

**Supplementary Table 8. Analysis results of G6PD metabolomics data using Metaboverse collapse reaction function.** A connection between dUMP and THF can be identified, but the reaction ranks lower, and no link between G6PD and the two can be established.

**Supplementary Table 9. The result of predicted paths ranking.** G6PD → dUMP

**Supplementary Table 10. The result of predicted paths ranking.** G6PD → AICAR

**Supplementary Table 11. The result of predicted paths ranking.** G6PD → FGAR.

**Supplementary Table 12. Statistics on the colorectal cancer cohort.** This table provides detailed demographic and clinical characteristics of the colorectal cancer cohort.

**Supplementary Table 13. MetaKG's predictions on the relationships between metabolites and colorectal cancer provide valuable insights.** Each metabolite is predicted to determine whether it has normal or abnormal regulation in connection with colorectal cancer.

**Supplementary Table 14. 235 aging related metabolites gathered from publications.**

**Supplementary Table 15. Comparisons between MetaKG and other metabolomic tools and databases.**

### AI-assisted editing

OpenAI Codex assisted with language editing, formatting and consistency checks of the existing manuscript. It was not used to generate experimental data or research results.
